# Impact of reference design on estimating SARS-CoV-2 lineage abundances from wastewater sequencing data

**DOI:** 10.1101/2023.06.02.543047

**Authors:** Eva Aßmann, Shelesh Agrawal, Laura Orschler, Sindy Böttcher, Susanne Lackner, Martin Hölzer

## Abstract

**Background:** Sequencing of SARS-CoV-2 RNA from wastewater samples has emerged as a valuable tool for detecting the presence and relative abundances of SARS-CoV-2 variants in a community. By analyzing the viral genetic material present in wastewater, public health officials can gain early insights into the spread of the virus and inform timely intervention measures. The construction of reference datasets from known SARS-CoV-2 lineages and their mutation profies has become state-of-the-art for assigning viral lineages and their relative abundances from wastewater sequencing data. However, the selection of reference sequences or mutations directly affects the predictive power.

**Results:** Here, we show the impact of a *mutation-* and *sequence-based* reference reconstruction for SARS-CoV-2 abundance estimation. We benchmark three data sets: 1) synthetic “spike-in” mixtures, 2) German samples from early 2021, mainly comprising Alpha, and 3) samples obtained from wastewater at an international airport in Germany from the end of 2021, including first signals of Omicron. The two approaches differ in sub-lineage detection, with the marker-*mutation-based* method, in particular, being challenged by the increasing number of mutations and lineages. However, the estimations of both approaches depend on selecting representative references and optimized parameter settings. By performing parameter escalation experiments, we demonstrate the effects of reference size and alternative allele frequency cutoffs for abundance estimation. We show how different parameter settings can lead to different results for our test data sets, and illustrate the effects of virus lineage composition of wastewater samples and references.

**Conclusions:** Here, we compare a *mutation-* and *sequence-based* reference construction and assignment for SARS-CoV-2 abundance estimation from wastewater samples. Our study highlights current computational challenges, focusing on the general reference design, which significantly and directly impacts abundance allocations. We illustrate advantages and disadvantages that may be relevant for further developments in the wastewater community and in the context of higher standardization.

## Background

Coronavirus disease 2019 (COVID-19), the highly contagious viral illness caused by severe acute respiratory syndrome coronavirus 2 (SARS-CoV-2), is the most consequential global health crisis since the era of the influenza pandemic of 1918. Since its discovery, SARS-CoV-2 has caused 763 million confirmed cases of COVID-19 (covid19.who.int, accessed April 19, 2023) and currently 3,000 SARS-CoV-2 lineages are defined by the *Pango* network (O’Toole et al., 2021; Rambaut et al., 2020) (https://github.com/cov-lineages/lineages-website/blob/master/_data/lineage_data.full.json, accessed April 19, 2023). Genome sequencing has played a central role during the COVID-19 pandemic in supporting public health agencies, monitoring emerging mutations in the SARS-CoV-2 genome, and advancing precision vaccinology and optimizing molecular tests (consortium, 2020; Robishaw et al., 2021; Oh et al., 2022). Massive sequencing of clinical samples has made it possible to monitor emerging variants, emphasizing temporal and spatial variation. With ongoing transmission, further mutations occur in the genome that are part of the viral evolutionary process and result in unique fingerprints.

Sequencing capacity, however, is limited, cannot be sustained over the long term for so many clinical samples, and only allows extrapolation based on a relatively small fraction of all infections occurring during the pandemic. In addition, with decreasing incidence numbers, sampling and sequencing efforts are decreasing, raising the need for representative, medium-scale, and sustainable surveillance systems (Oh et al., 2022) or other approaches. From January 1, 2020 until April 19, 2023, 931,260 genome sequences of COVID-19-positive clinical samples from Germany have been uploaded to the international GISAID platform (Shu and McCauley, 2017), representing a proportion of 2.426 % out of a total of 38,388,247 reported SARS-CoV-2 cases in Germany (COVID-19 Dashboard Germany, accessed April 19, 2023). In Germany and other countries, complete detection and sequencing of all positive cases were impossible due to the high infection numbers. However, wastewater-based epidemiology (WBE) has shown the potential to get a much broader snapshot of the SARS-CoV-2 variant circulation at a community level (Jahn et al., 2022; Smyth et al., 2022; Agrawal et al., 2022b; Peccia et al., 2020; Nemudryi et al., 2020; Hoar et al., 2022). Integrating genome sequencing with WBE can provide information on circulating SARS-CoV-2 variants in a region (Amman et al., 2022). The sequencing methods commonly used in WBE are similar to the ones used for clinical samples, using a general strategy that employs the sequencing of the whole genome via amplification of small, specific regions of the SARS-CoV-2 genome, i.e., targeted sequencing of amplicons via pre-defined primer sequences (Gregory et al., 2021; Barbé et al., 2022; Smyth et al., 2022; Agrawal et al., 2022a; Nemudryi et al., 2020). Targeted sequencing can achieve a high degree of coverage of informative regions of the genome and, most importantly, reveal to some extent which polymorphisms are linked, making it possible to track SARS-CoV-2 variants of concern (VOCs) and other variants.

A particular challenge in performing sequencing of SARS-CoV-2 from wastewater samples concerns the viral RNA present in many individual fragments rather than complete viral genomes. In addition, these fragments come from the excretions of many infected individuals, making it challenging, if not impossible, to reconstruct individual genomes using bioinformatic approaches like the ones developed for clinical samples of individual patients. In need of computational approaches to analyze mixed wastewater samples, several groups developed similar tools for quality control, sequencing data analysis, and SARS-CoV-2 lineage abundance estimation (Karthikeyan et al., 2022; Pechlivanis et al., 2022; Valieris et al., 2022; Ellmen et al., 2021; Amman et al., 2022; Barker et al., 2021; Schumann et al., 2022; Gregory et al., 2021; Jahn et al., 2022; Gafurov et al., 2022; Posada-Céspedes et al., 2021; Baaijens et al., 2022; Korobeynikov, 2022; Kayikcioglu et al., 2023), see ***Table 1***. Most approaches focus on detecting pre-defined characteristic marker mutations in the sequenced reads and utilize this information for abundance estimation. Common to all these tools is that they require a reference set of either signature marker mutations (hereafter called *mutation-based*) or complete genome sequences (hereafter called *sequence-based*) from which characteristic mutation profiles or kmers (short sub-sequences of length k) are derived. Kayikcioglu *et al*. compared the performance of 1ve selected approaches for SARS-CoV-2 lineage abundance estimation on simulated and publicly available mixed population samples (Kayikcioglu et al., 2023). They found that Kallisto (Bray et al., 2016), as first suggested by Baaijens, Zulli, and Ott *et al*. (Baaijens et al., 2022), followed by Freyja (Karthikeyan et al., 2022), achieved most accurate estimations.

**Table 1.** Collection of tools available for sequencing data analysis in WBE and SARS-CoV-2 lineage proportion estimation. We distinguish the tools roughly based on their approach to define a reference set into those using predefined marker mutations and those relying on full genome sequences or both. The two implementations we selected for reference construction and our comparison are indicated in bold. Please note that C-WAP (Kayikcioglu et al., 2023) wraps multiple approaches while also including a new *mutation-based* tool, LINDEC.

| ***mutation-based*** |  |  |
| --- | --- | --- |
| Tool | Citation | Code |
| **MAMUSS** | (Agrawal et al., 2022a) | github.com/lifehashopes/MAMUSS |
| Freyja | (Karthikeyan et al., 2022) | github.com/andersen-lab/Freyja |
| Lineagespot | (Pechlivanis et al., 2022) | github.com/nikopech/lineagespot |
| LCS | (Valieris et al., 2022) | github.com/rvalieris/LCS |
| Alcov | (Ellmen et al., 2021) | github.com/Ellmen/alcov |
| VaQuERo | (Amman et al., 2022) | github.com/fabou-uobaf/VaQuERo |
| MMMVI | (Barker et al., 2021) | github.com/dorbarker/voc-identify |
| PiGx | (Schumann et al., 2022) | github.com/BIMSBbioinfo/pigx_sars-cov-2 |
| SAMRefiner | (Gregory et al., 2021) | github.com/degregory/SAM_Refiner |
| COJAC | (Jahn et al., 2022) | github.com/cbg-ethz/cojac |
| wastewaterSPAdes | (Korobeynikov, 2022) | – |
| gromstole | – | github.com/PoonLab/gromstole |
| CovMix | – | github.com/chrisquince/covmix |
| ***sequence-based*** |  |  |
| Tool | Citation | Code |
| **VLQ-nf** | this study | github.com/rki-mf1/VLQ-nf |
| VLQ | (Baaijens et al., 2022) | github.com/baymlab/wastewater_analysis |
| VirPool | (Gafurov et al., 2022) | github.com/fmfi-compbio/virpool |
| V-pipe SC2 | (Posada-Céspedes et al., 2021) | cbg-ethz.github.io/V-pipe/sars-cov-2 |
| ***mutation-based & sequence-based*** |  |  |
| Tool | Citation | Code |
| **C-WAP** | (Kayikcioglu et al., 2023) | github.com/CFSAN-Biostatistics/C-WAP |

In a *mutation-based* approach, to estimate the proportion of specific SARS-CoV-2 variants present in a mixed sample, mutations or combinations of mutations characteristic or unique for these variants based on clinical samples can be compared with the mutations detectable in the sample. In principle, and as implemented in a previously used approach (Agrawal et al., 2022a) (which we refer to here as MAMUSS, ***Table 1***), the occurrence of mutations can be represented by the value of the relative abundance of a VOC or other viral variant. First, the frequency of occurrence of each mutation is calculated from the multiplication of the reads and the allele frequency. The relative abundance describes the percentage ratio of the sum of the read abundance of the characteristic mutations of a SARS-CoV-2 virus variant and the sum of the read abundance of all mutations found in a sample. Accordingly, only the previously selected virus variants and signature mutations that form the reference set are evaluated and others that may occur in the sample are ignored. Another prominent *mutation-based* approach is implemented in the tool Freyja (Karthikeyan et al., 2022). Freyja solves the de-mixing problem to recover relative lineage abundances from mixed SARS-CoV-2 samples using lineage-determining mutational “barcodes” derived from the UShER global phylogenetic tree (Turakhia et al., 2021). Using mutation abundances and sequencing depth measurements at each position in the genome, Freyja estimates the abundance of lineages in the sample.

As a different methodological approach to reconstruct a reference, the full genome sequence information can be used to automatically select appropriate features (e.g., signature mutations, kmers) and to use them to evaluate the proportions of SARS-CoV-2 variants in wastewater samples instead of a pre-selected set of marker mutations (*sequence-based*) (Baaijens et al., 2022; Gafurov et al., 2022; Posada-Céspedes et al., 2021) ***Table 1***. Again, information derived from sequencing of clinical samples and their lineage annotation are used to generate a representative reference data set that can be then searched via established (pseudo)-alignment methods such as Kallisto (Bray et al., 2016) as suggested by Baaijens, Zulli, and Ott *et al*. in their VQL tool (Baaijens et al., 2022).

In this study, we specifically investigated the impact of reference composition and construction on assigning relative abundances of SARS-CoV-2 lineages from wastewater sequencing data. As mentioned, various tools have been developed over the course of the pandemic (***Table 1***) and they all have different facets in calculating relative abundances. However, they all have in common that some reference data set needs to be defined, which derives information on lineages and mutations from existing genomics data. We specifically chose two different approaches representing a *mutation-based* method (MAMUSS) and *sequence-based* method (VLQ). The two approaches distinguish mainly by the input data set used for the reference set design and subsequent lineage assignment (***Figure 1***). Either a selection of lineage-defining marker mutations (*mutation-based*) or full SARS-CoV-2 genome sequences (*sequence-based*) are used to reconstruct a reference base for lineage assignment and abundance estimation. Here, we compare exemplary implementations of both general approaches. MAMUSS, as previously applied in (Agrawal et al., 2022a), implements a representative basic workflow for the *mutation-based* approach focusing on unique marker mutations. For the *sequence-based* approach, we use pseudo-alignments via Kallisto (Bray et al., 2016) as proposed initially by (Baaijens et al., 2022) and their VQL tool. Based on their idea and scripts, we implemented a slightly modified version of VLQ in a Nextflow (Di Tommaso et al., 2017) pipeline that we call VLQ-nf. We chose our *sequence-based* method to be based on VLQ because of it reusing Kallisto as an established tool in transcript quantification (Bray et al., 2016). A major benefit of implementing the representative methods was the complete control over code, parameters, and inputs, which allowed us to understand better, compare, and interpret the results of our benchmark study and the effects on the reference design.

**Figure 1.**
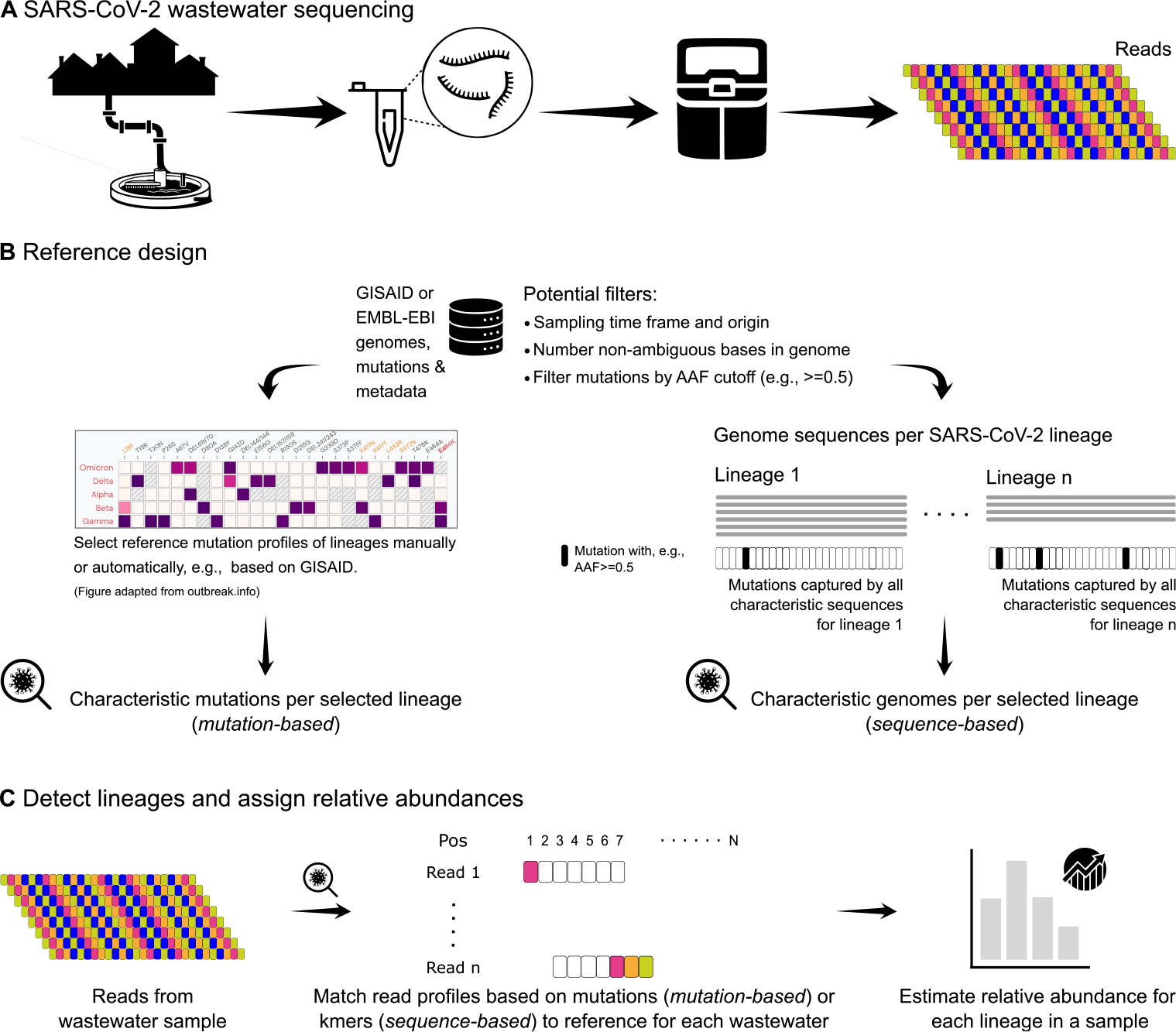
Schematic overview of reference design and lineage abundance estimation from SARS-CoV-2 wastewater sequencing data. (A) Wastewater samples are collected from sewer influent, for example. RNA is extracted and, in the context of SARS-CoV-2, usually amplified as cDNA using established primer schemes and then sequenced to obtain short snippets of viral RNA (*reads*). (B) Current methods (***Table 1*** for lineage assignment and abundance estimation need a reference data set, usually constructed from genomes and mutations derived from clinical sequencing and patient samples. Here, we distinguish two general approaches to design the reference, where either marker mutations are pre-selected (*mutation-based*) or full-genome sequences are selected (*sequence-based*). (C) The data analysis part may differ significantly depending on the implementation. However, all tools attempt to assign known lineages and estimate their frequency in the mixed sample based on mutations that can be detected in the reads. Our study uses MAMUSS as an exemplary *mutation-based* approach based on a two-indicator classification and pre-selected marker mutations characteristic for certain lineages (Agrawal et al., 2022a). For the *sequence-based* approach, we use a Nextflow implementation (VQL-nf) of the slightly adjusted VLQ pipeline as proposed by Baaijens, Zulli, and Ott *et al*. and which is based on the tool Kallisto (Baaijens et al., 2022). AAF – Alternative Allele Frequency, used as a cutoff to define a mutation as a feature.

We tested both MAMUSS as a *mutation-based* reference representative and VLQ-nf as a *sequence-based* reference representative on three data sets: 1) a synthetic scenario of “spike-in” mixture samples, 2) samples from Germany from a European wastewater study from early 2021, mainly comprising the VOC Alpha (Agrawal et al., 2022b), and 3) a sample obtained from wastewater sequencing at the international airport in Frankfurt am Main, Germany from the end of 2021, including 1rst signals of the VOC Omicron (Agrawal et al., 2022a).

We show that both the *mutation-based* and *sequence-based* approach can reflect the proportions of SARS-CoV-2 lineages in the different samples but also comprise differences in resolution and the detection of similar sub-lineages depending on the reference set. Both approaches also show advantages and disadvantages when it comes to the selection of signature marker mutations and genome sequences, respectively. For the *mutation-based* approach as implemented in MAMUSS, it became more and more challenging to select (sub-)lineage-defining marker mutations that provide robust assignments in the context of the increasing diversity of SARS-CoV-2 lineages.

## Data Description

### Data collection and benchmark scenarios

We selected three wastewater data sets for our comparison to cover 1) a synthetic scenario of “spike-in” mixture samples (*Standards*; n=16 samples), 2) real samples from early 2021 from a large European study and collected in Germany (Agrawal et al., 2022b), mainly comprising the VOC Alpha (*Pan-EU-GER*; n=7 samples), and 3) one sample from the end of 2021 including first signals of the VOC Omicron obtained from wastewater at the international airport in Frankfurt am Main, Germany (*FFM-Airport*; n=1 sample) (Agrawal et al., 2022a). The *Standards* comprise RNA from 10 SARS-CoV-2 variants (including the original Wuhan-Hu-1 A.1 lineage), which were mixed in different proportions to generate 16 samples for library preparation and sequencing via Ion Torrent (***Table 2***). Please note that no real wastewater was used to construct the *Standards* (see Methods). Within the Pan-EU WBE study, high-quality sequencing data was produced for SARS-CoV-2 wastewater samples across 20 European countries, including 54 municipalities (Agrawal et al., 2022b). We selected the seven German samples from this study (SRX11122519 and SRX11122521–SRX11122526; *Pan-EU-GER*) for our benchmark, which were sampled in March 2021 and mainly cover the rise of the VOC Alpha during that time. Lastly, we obtained one sample (SRR17258654) from wastewater sampling in November 2021 at the international airport in Frankfurt am Main (*FFM-Airport*) and where we found first signals and low proportions of the VOC Omicron arriving during that time in Germany (Agrawal et al., 2022a).

**Table 2.** Composition of synthetic mixture “spike-in” *Standards*. Here we show the proportions of which different SARS-CoV-2 lineages were mixed to generate a collection of artificial samples for our benchmark. For example, the sample Mix_01 comprises 25 % original Wuhan-Hu-1 A.1 and 75 % Alpha B.1.1.7 (0.25*_org_* − 0.75*_alpha_*). All samples were sequenced with Ion Torrent and raw data is available under the BioProject number PRJNA912560 in the National Center for Biotechnology Information (NCBI) Seuqence Read Archive (SRA). Please note that no real wastewater was used to construct these synthetic mixtures because we wanted to reduce any side effects for our *gold standard*.

| Sample ID | Composition |
| --- | --- |
| Mix_01 | $0.25_{org} - 0.75_{alpha}$ |
| Mix_02 | $0.25_{org} - 0.25_{beta} - 0.5_{alpha}$ |
| Mix_03 | $0.25_{alpha} - 0.25_{beta} - 0.25_{gamma} - 0.25_{org}$ |
| Mix_04 | $0.5_{org} - 0.5_{iota}$ |
| Mix_05 | $0.25_{org} - 0.25_{iota} - 0.5_{omiba2}$ |
| Mix_06 | $0.25_{alpha} - 0.25_{iota} - 0.25_{omiBA1} - 0.25_{omiBA2}$ |
| Mix_07 | $0.5_{omiBA1} - 0.5_{omiBA2}$ |
| Mix_08 | $0.25_{org} - 0.25_{alpha} - 0.25_{omiBA1} - 0.25_{omiBA2}$ |
| Mix_09 | $0.5_{deltaAY1} - 0.5_{deltaAY2}$ |
| Mix_10 | $0.25_{deltaAY1} - 0.25_{deltaAY2} - 0.5_{delta}$ |
| Mix_11 | $0.25_{deltaAY1} - 0.25_{deltaAY2} - 0.5_{omiBA1}$ |
| Mix_12 | $0.25_{deltaAY1} - 0.25_{deltaAY2} - 0.25_{omiBA1} - 0.25_{omiBA2}$ |
| Mix_13 | $0.25_{deltaAY1} - 0.25_{deltaAY2} - 0.25_{omiBA1} - 0.25_{omiBA2}$ |
| Mix_14 | $0.5_{delta} - 0.25_{omiBA1} - 0.25_{omiBA2}$ |
| Mix_15 | $0.25_{deltaAY1} - 0.25_{deltaAY2} - 0.25_{omiBA1} - 0.25_{omiBA2}$ |
| Mix_16 | $0.25_{alpha} - 0.25_{delta} - 0.25_{omiBA1} - 0.25_{omiBA2}$ |

*org* – Wuhan-Hu-1 A.1; *alpha* – Alpha B.1.1.7; *beta* – Beta B.1.351; *gamma* – Gamma P.1; *iota* – Iota B.1.526; *delta* – Delta B.1.617.2; *deltaAY1* – Delta AY.1; *deltaAY2* – Delta AY.2; *omiBA1* – Omicron BA.1; *omiBA2* – Omicron BA.2

### Approaches for SARS-CoV-2 reference reconstruction and lineage abundance estimation from wastewater

We compare two approaches based on different types of reference construction to assign lineages to SARS-CoV-2 wastewater sequencing data for downstream abundance estimation. Our *mutation-based* approach, MAMUSS, defines lineage-defining marker mutations, while our implementation of a *sequence-based* approach, VLQ-nf, utilizes full SARS-CoV-2 genome sequences to build a reference database. We compared both approaches on three wastewater sequencing data sets described above. For MAMUSS, the variant surveillance database from GISAID (https://www.gisaid.org) (Shu and McCauley, 2017) is used to reconstruct reference mutation profiles for each virus variant. Mutations that have been often reported for each virus variant are considered for reference profiles. SARS-CoV-2 variants in wastewater samples are determined by comparing the mutation profiles, generated using a variant caller, with the constructed reference mutation profiles. The abundance estimation is based on the read depth and allele frequency of each mutation detected in a wastewater sample. For the *sequence-based* approach, we implemented an adjusted version of the VQL scripts by Baaijens, Zulli, and Ott *et al*. (Baaijens et al., 2022), re-using the pseudo-aligner Kallisto (Bray et al., 2016) as a Nextflow pipeline (VLQ-nf). This method reconstructs a reference index from GISAID SARS-CoV-2 whole-genome sequences (Shu and McCauley, 2017). The exact lineages and genome sequences considered for reference reconstruction are selected based on filters controlling the represented temporal and geographical range, as well as the level of genomic variation among the included lineages. SARS-CoV-2 sequencing data is then pseudo-aligned against the reference index to assign lineages. The abundance estimation is performed by an Expectation-Maximization algorithm.

### Data availability

All used raw sequencing data files for the *Pan-EU-GER* and *FFM-Airport* data sets were uploaded to ENA in the context of their original publications (Agrawal et al., 2022b; Agrawal et al., 2022a). The sequencing data for the *Standards* benchmark are available under the NCBI BioProject number PRJNA912560. Further intermediate results and data files (reference sequences, constructed indices) can be found at osf.io/upbqj for full reproducibility of our analyses. The code for our Nextflow implementation based on the proposed method and original code by Baaijens, Zulli, and Ott *et al*. (Baaijens et al., 2022) is freely available at https://github.com/rki-mf1/VLQ-nf. Code for the *mutation-based* approach MAMUSS is freely available at https://github.com/lifehashopes/MAMUSS.

## Results

### Both the ***mutation-based*** and ***sequence-based*** approaches yield similar SARS-CoV-2 lineage proportions for mixed ***Standard*** samples but differ on sub-lineage level

We analyzed our *Standards* data set (***Table 2***) using the *sequence-based* approach implemented in VLQ-nf and an implementation of a *mutation-based* approach, MAMUSS (***Table 1***). Given ground truth knowledge, we assessed the qualitative and quantitative performance of both methods yielding controlled insights into the strengths and limitations of each approach.

VLQ-nf detected all correct spike-in lineages across all samples. The output for every sample showed, however, a certain amount of false positive predictions comprising lineages that are part of our reference set but not used as spike-ins (***Figure 2***). We observed the most consistent false positive estimations for Gamma (P.1) with up to 1.61 % abundance across all samples. In contrast, MAMUSS did not detect all spike-in lineages, but also showed more robust results in quantifying fewer false positives in the samples (***Figure 2***).

**Figure 2.**
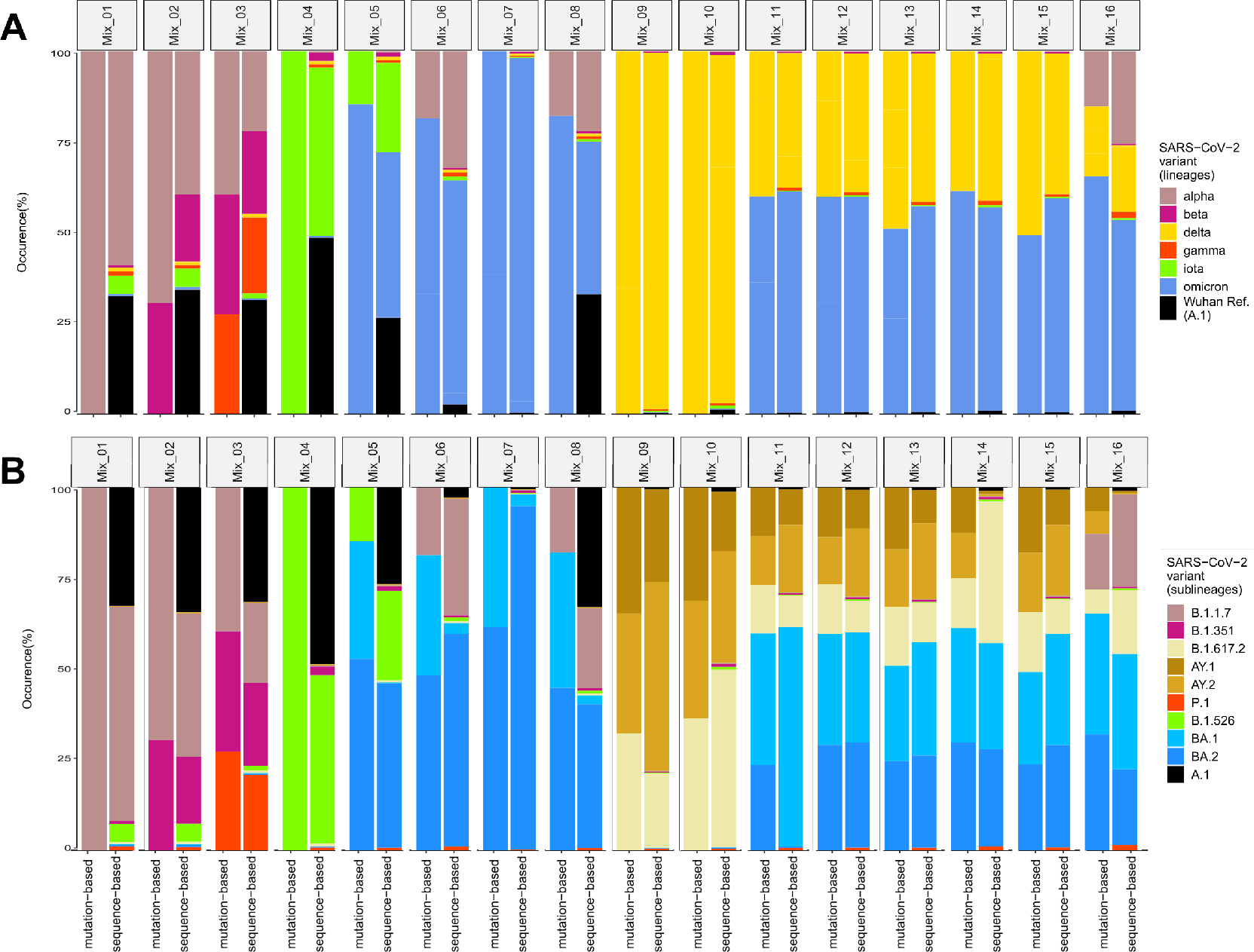
Comparison of the occurrence of pre-defined mixtures of SARS-CoV-2 variants (*Standards*) (A) at Pangolin parent lineage level and (B) at Pangolin sub-lineage resolution based on the *sequence-based* (VLQ-nf) and *mutation-based* (MAMUSS) approach.

When comparing false detection and over- or underestimation for both approaches, we partly observed similar patterns among specific groups of lineages: The *mutation-based* approach showed a bias in samples comprising A.1 towards not being able to detect A.1 and instead detecting false positives of B.1.1.7 or BA.1. In sample Mix_06, the *mutation-based* approach could not detect Iota (B.1.526) but falsely detected BA.1. Similarly, the *sequence-based* approach showed varying patterns of over- and underestimations among B.1.526 and B.1.1.7 (see Mix_01, Mix_02, and Mix_06).

Furthermore, both approaches showed distinct patterns of false estimation among B.1.617.2 (Delta) and its sub-lineages AY.1 and AY.2. In samples containing no Delta and only Delta sub-lineages, both approaches falsely detected Delta while underestimating AY.1 or AY.2. In samples containing only Delta and no Delta sub-lineages, MAMUSS falsely detected AY.1 and AY.2, while underestimating Delta. In samples containing both Delta and Delta sub-lineages, VLQ-nf overestimated Delta and underestimated AY.1, while MAMUSS overestimated Delta sub-lineages and underestimated Delta.

Both approaches estimated BA.1 and BA.2 without distinct conflicts among each other. We observed slight over- or underestimation in the abundance of Omicron lineages to co-occur with underestimation of Delta sub-lineages in samples Mix_10–16.

Finally, we found both approaches to match the ground truth proportions of the *Standards* samples well on the parent lineage level. On the sub-lineage level, we found the false negative detection of B.1.526 in sample Mix_06 and the quantification conflicts among Delta (sub-)lineages to be the most prominent conflicts among both approaches. For the *mutation-based* approach, we found the false negative detection of A.1 to be the second most prominent shortcoming observed in this experiment.

### VLQ-nf detects Alpha sub-lineages while MAMUSS ***1***nds distinctly larger abundances for rising lineages Beta, Gamma, and Delta in the ***Pan-EU-GER*** data

We analyzed German samples from the Pan-EU study (Agrawal et al., 2022b) using both approaches to assess their performance on real wastewater sequencing data. In the lack of ground truth knowledge, we evaluated both approaches by relating the lineage predictions and quantification to the pandemic background in Germany based on data from clinical sampling strategies. Moreover, we performed experiments on real data to evaluate the potential benefits of wastewater-based surveillance compared to clinically-based data.

According to global surveillance projects based on clinical genomic sequence data such as outbreak.info (Gangavarapu et al., 2023) (accessed June 03, 2022) and Nextstrain (Hadfield et al., 2018) (accessed June 03, 2022), the pandemic situation in Europe from February until April 2021 was mainly dominated by the SARS-CoV-2 lineages Alpha, Beta, cases of B.1.177 and sub-lineages, B.1.258 and sub-lineages, and B.1.160 (Supplementary ***Figure S1***). The pandemic situation in Germany at that time was mainly dominated by Alpha, B.1.177.86, B.1.177.81, Beta, B.1.258, B.1.177, and B.1.160. According to GISAID submissions during that time, approximately the same lineages and multiple other low-abundant global and European sub-lineages were reported from clinical sampling strategies. Here we focused the comparison on the lineages Alpha (B.1.1.7), Beta (B.1.351), Gamma (P.1), Delta (B.1.617.2), and the respective sub-lineages, as those were or became the dominant lineages around the time of wastewater sampling in Germany in the context of the Pan-EU project (Agrawal et al., 2022b).

With VLQ-nf, we quantified the lineage and sub-lineage level. In comparison, MAMUSS predicted lineage abundances only at the parent level (***Figure 3***). Both approaches predicted Alpha (sub-)lineages to be the most abundant lineages in the data set. Specifically, the *sequence-based* approach found Alpha sub-lineages Q.1 and Q.7 to be the most abundant. Yet, those Alpha sub-lineages were not reported amongst the most frequent cases based on clinical sampling strategies (see Supplementary ***Figure S1***). We also detected Beta, Gamma, and Delta (sub-)lineages at abundances below 1 %, which are not visible at the scale of ***Figure 3***. In contrast, we found distinctly larger abundances of Beta, Gamma, and Delta in the samples using MAMUSS.

**Figure 3.**
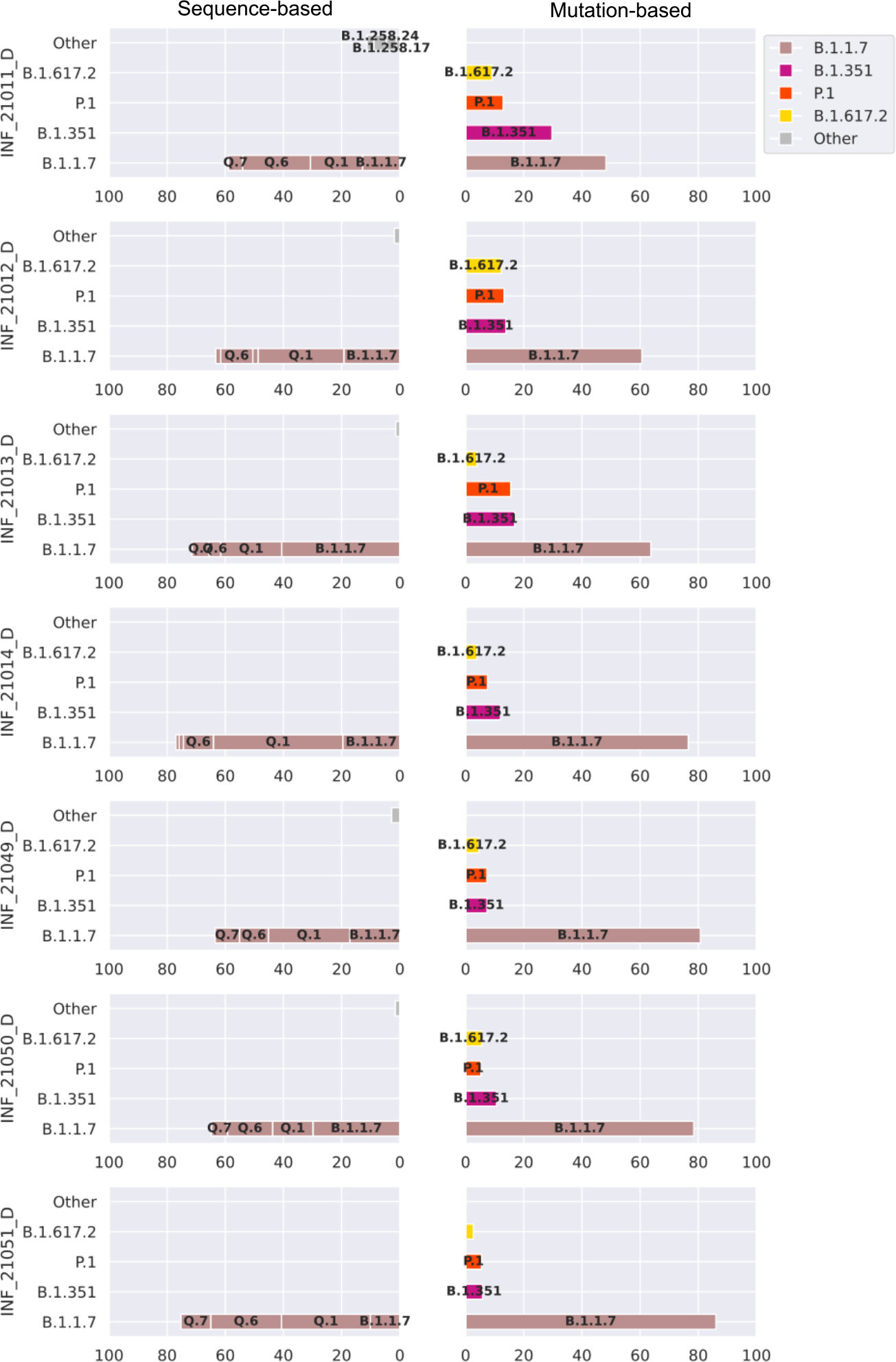
Comparison of the results for the *Pan-EU-GER* analysis using VLQ-nf (*sequence-based*, left) versus MAMUSS (*mutation-based*, right). Abundance predictions are plotted above a cutoff of 1 % abundance and labeled at a threshold of 3 % abundance. VLQ-nf detected abundances for B.1.617.2, P.1, and B.1.351 sub-lineages below 1 %, which is not visible at the scale of this 1gure.

### ***Mutation***- and ***sequence-based*** approaches recover a similar Omicron proportion from an early airport wastewater sample

We used both approaches to analyze a real wastewater sequencing sample (SRR17258654, *FFM- Airport*) (Agrawal et al., 2022a). We compared lineage predictions and quantification against the pandemic background in Europe and South Africa at the time of wastewater sampling. We evaluated both approaches in terms of their ability to detect (sub-)lineages at low abundances, specifically to detect low abundant signals of Omicron.

The pandemic situation in Europe and South Africa from October to December 2021 was dominated by Delta sub-lineages and increasing incidences of Omicron and its sub-lineages according to outbreak.info (Gangavarapu et al., 2023) and nextstrain.org (Hadfield et al., 2018) (Supplementary ***Figure S2***). According to GISAID submissions, mostly Delta sub-lineages and a few cases of Omicron and other minor global sub-lineages were reported based on clinical sampling strategies.

With VLQ-nf, we detected many Delta sub-lineages at abundances ranging from less than 1 % to around 8 % that in sum contribute over 93 % abundance in the wastewater sample (***Figure 4***). We observed BA.1 with 1.44 % and some other lineages and sub-lineages with abundances of less than 1 % (“Other”) in that sample. We found all lineage quantification of less than 1 % to aggregate to around 48 % abundance in total.

**Figure 4.**
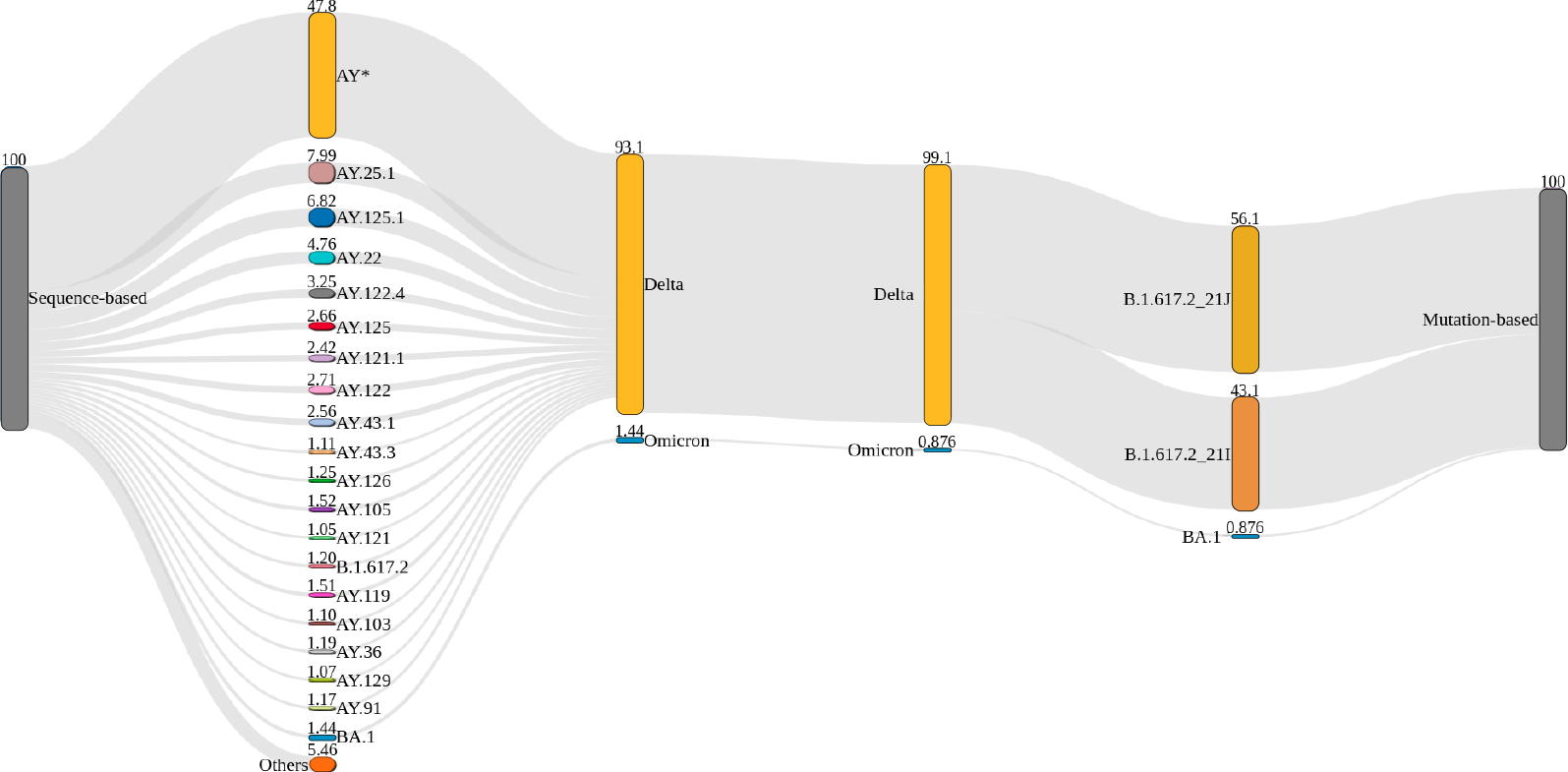
Sankey plot comparing the detected lineage proportions for the *sequence-based* approach (VQL-nf, left) and the *mutation-based* approach (MAMUSS, right) for one airport wastewater sample (SRR17258654) (Agrawal et al., 2022a). Both approaches detect a similar amount of Delta and Omicron (BA.1) in the sample, while VQL-nf can achieve a higher sub-lineage resolution (AY lineages) based on the full genome information in the reconstructed reference index and utilizing pseudo-alignments. MAMUSS can, as configured for this analysis and based on the limited reference set, distinguish between two slightly different B.1.617.2 clades as defined by Nextstrain. For the *sequence-based* approach, only lineages with a proportion of at least 1 % are shown and all other AY-sub-lineages are pooled in AY* and all other lineages in ”Others”.

We observed a similar lineage abundance profile with MAMUSS. We found that most abundance consists of two approximately equally abundant Delta sub-lineages. We detected a small proportion close to 1 % of Omicron. Compared to VLQ-nf, we did not 1nd any low abundant quantification for other (sub-)lineages, explained by the smaller reference data set only composed of a particular collection of marker mutations.

We found that the estimated abundance profiles of lineages from both approaches matched well with the pandemic background in Europe and South Africa at the time of wastewater sampling. However, when considering abundance estimations of the *sequence-based* approach at the sub- lineage level, we discovered differences regarding the most abundantly predicted Delta sub-lineages compared to the more prominent Delta sub-lineages derived from clinical sampling strategies in European and South African GISAID submissions. The *sequence-based* approach predicted AY.25.1, AY.125.1, AY.122.4, AY.121, and AY.43.1 to be most abundant in the analyzed sample. In contrast, GISAID submissions showed AY.4, AY.43, AY.122, AY.4.2, AY.126, AY.4.2.2, and AY.98 as the most frequent Delta sub-lineages in Europe during that time. Additionally, we found AY.45, AY.32, AY.91, AY.116, AY.122, AY.6, and AY.46 to be the highest reported Delta sub-lineages in South Africa. While our predictions do not match the clinically reported frequencies, some of our predictions belong to the same lineage family as the most frequently reported lineages from clinical sampling, e.g., AY.43.1 is a sub-lineage of AY.43, AY.122.4 is a sub-lineage of AY.122, and AY.125.1 is a sub-lineage of AY.125 which we found among the twenty most frequently reported lineages in Europe using VLQ-nf.

### Alternative allele frequency and size of reference database impact the ***sequence- based*** method, but the effects also depend on lineage composition in the sample

To better understand the impact of specific parameters on the performance of the *sequence-based* method, we performed parameter escalation experiments on the *Standards* benchmark set as well as the *PanEU-Ger* and *FFM-Airport* data sets. Due to the similar findings for all three data sets, here we only present the results based on the *Standards* and refer to the results of the *PanEU-Ger* and *FFM-Airport* data sets in the Supplement. We investigated the impact of reference construction parameters on lineage proportion estimation and aimed at uncovering the potential bias of the pseudo-alignment implemented in the *sequence-based* method. Specifically, we focused on the AAF threshold and the maximum number of sequences per lineage. The AAF threshold defines the minimum alternative allele frequency for a mutation to be considered characteristic of a lineage. First, genome sequences are added as lineage references so that each mutation that exceeds the AAF threshold is detected at least once by as few sequences as possible. Next, additional genomes are added until the maximum number of sequences per lineage is reached. Thus, the AAF threshold controls the level of genomic variation captured for each lineage and the maximum number of sequences per lineage controls the reference size.

### Standards

Across most *Standards* samples and experiments, VLQ-nf detected all spike-in lineages and predicted reasonable estimates (***Figure 5***). However, we consistently observed low abundant false positive hits in all of our mixed samples, comprising lineages that are part of the reference index but not used as spike-ins. We found the most prominent false positive detection to be Gamma. We observed similar patterns of false positive detection and false estimation among specific groups of lineages across all parameter settings: For the first eight samples Mix_01 to Mix_08, most cases of false estimation of spike-in lineage abundances occurred alongside false positives or negatives of B.1.526 and false positives of BA.1. For the samples Mix_09 to Mix_16, we observed most detection con2icts to involve ambiguities among Delta and its sub-lineages AY.1 and AY.2.

**Figure 5.**
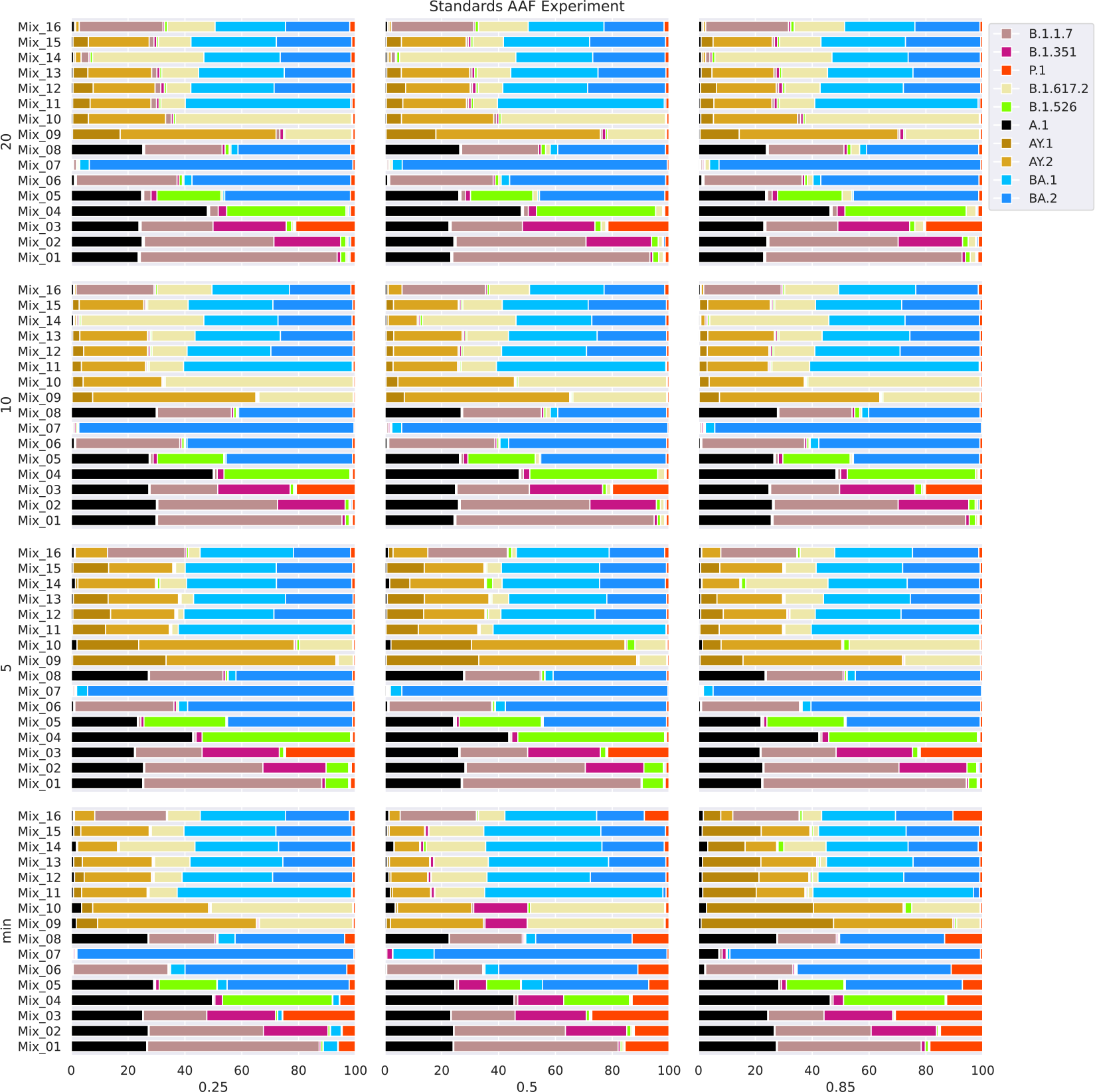
Results for the parameter escalation experiments on the *Standards* samples using the *sequence-based* method (VLQ-nf). We analyzed the *Standards* with different parameterizations for reference construction (x-axis: increasing AAF threshold, y-axis: increasing maximum number of sequences per lineage). VLQ-nf using pseudo-alignments detected all lineages and estimated abundance profiles well across most samples and parameter settings. However, we also observed prominent detection ambiguities among Delta and its sub-lineages and found consistently low abundant false positives for specific groups of lineages. Continuously increasing or decreasing parameter settings caused heterogeneous changes in the estimated abundance proportions across samples. The *sequence-based* method showed to perform better when using a reference set larger than the minimum reference size. Still, we found noise levels to increase distinctly when using the maximum reference size among the considered settings.

We found that the detection and quantification performance of the *sequence-based* method via VLQ-nf changed with varying parameter settings. Specifically, we found those changes to vary across samples and observed them not to behave identically with consistent parameter changes. For example, at the minimum reference size (Supplementary ***Table S1***), we observed abundance predictions for samples Mix_09 and Mix_11–16 to first improve with an increasing AAF threshold. However, with a further increasing AAF threshold, we observed more false estimations of Delta sub-lineages. Furthermore, although Mix_10 shares most of its spike-in lineages with Mix_09, the performance of abundance estimations for sample Mix_10 1rst decreased and then improved again when increasing the AAF threshold.

We made a similar observation for the maximum number of sequences per lineage. With an AAF threshold of 0.5, the abundance estimates for Mix_01 improved with increasing reference size, while we found them to deteriorate for Mix_09 which includes a distinctly different sample composition. Overall, we found lineage abundance estimations to become slightly more robust across varying AAF thresholds with increasing reference size. This is reflected best in the abundance profiles for samples Mix_09–Mix_15 when looking at the proportional changes across increasing AAF settings for the minimum reference size throughout the reference with 10 sequences per lineage.

Finally, we found that the AAF threshold and the reference size affect the performance of the *sequence-based* method. Although we did not observe a clear and consistent pattern of impact, we found that the effects of varying parameter settings may depend on the sample composition. Specifically, we observed the strongest impact of parameter changes for samples containing lineages with a higher degree of shared genomic similarity. Also, we found the AAF threshold to affect estimates slightly more than the reference size. We detected similar results for the *PanEU-Ger* and *FFM-Airport* data sets. We provide details for these two data sets in the Supplement (see ***Figure S3*** and ***Figure S4***).

### Final choice of parameters for benchmark reference construction

Within the scope of the parameter escalation experiments described here, we wanted to determine parameters with a good prediction performance without manipulating the benchmark in favor of the *sequence-based* approach (VQL-nf). Finally, based on our parameter testing and the three different data sets, we chose an AAF threshold of 0.25 and a reference size of at most 5 sequences per lineage. By that, we limit the size of the reference data set and still allow reasonable detection and quantification results across all three benchmark data sets, while at the same time keeping computational resources moderate.

## Discussion

It is apparent that the composition of the reference used must have a large impact on the deter- mination of relative SARS-CoV-2 abundances in wastewater sequence data. Especially given the dynamic and constantly updated SARS-CoV-2 lineage definitions (Rambaut et al., 2020), the refer- ence genome sequences and the signature mutations derived from them also change frequently.

Of course, the various tools (***Table 1***) and their parameters developed for estimating the relative abundance of lineages from wastewater sequencing data also have an impact. Here, however, we have specifically focused on the effects of the reference design.

We selected two general approaches for the design of reference data sets and estimation of SARS-CoV-2 lineage proportions from wastewater sequencing samples (***Figure 1***). On the one hand, selected marker mutations that are characteristic for certain SARS-CoV-2 lineages can be used for annotation and lineage proportion estimation (*mutation-based*, MAMUSS). Here, the read sequences derived from a wastewater sample are mapped against a reference genome from which differences (mutations) are detected and compared against the selected marker mutations. On the other hand, full SARS-CoV-2 genome sequences can be used to create a reference index without prior collection of specific mutations (*sequence-based*, VLQ-nf). Here, the problem of selecting appropriate marker mutations is shifted to the selection of representative lineages from which then features for the classification task are derived. An exemplary implementation of this approach based on the pseudo-aligner Kallisto (Bray et al., 2016) was recently proposed by Baaijens, Zulli, and Ott *et al*. (Baaijens et al., 2022). Based on their work, we developed a Nextflow pipeline for higher automation and reproducibility and the detection of SARS-CoV-2 lineage proportions from wastewater data using pseudo-alignments (VLQ-nf). In this approach, a selection of whole-genome SARS-CoV-2 sequences (target reference set) and the reads (query) are composed into kmers which are then eZciently compared to quantify lineage abundances, similar to quantifying gene expression in an RNA-Seq study.

To benchmark reference designs from both methods (*mutation-based* via MAMUSS, *sequence-based* via VLQ-nf), we selected three test scenarios: 1) a spike-in experiment with different SARS-CoV-2 lineage mixes, 2) samples obtained for Germany from a Pan-EU wastewater study, and 3) a wastewater sample from a German airport during the time when Omicron emerged.

In general, both approaches were able to detect SARS-CoV-2 lineage abundances from our test cases. The most remarkable difference was in the number of detected sub-lineages which also directly correlates with the reference design. VLQ-nf generally detected a larger diversity of sub- lineages in comparison to MAMUSS, which can be explained by the underlying reference indices. For the *mutation-based* approach and the implementation we used, it got increasingly diZcult to select a representative set of marker mutations. In contrast, the *sequence-based* approach as suggested by Baaijens, Zulli, and Ott *et al*. (Baaijens et al., 2022) can build a reference index on a large collection of SARS-CoV-2 full genome sequences and thus, potentially, better reflect diversity on sub-lineage levels. However, we also observed a certain amount of *noise* in the pseudo-alignment results causing potential false-positive hits in our test data sets. Other approaches, like Freyja (Karthikeyan et al., 2022), partly tackle this problem by deriving signature mutation profiles automatically, for example using the whole phylogenetic diversity of current SARS-CoV-2 sequences reflected in an UShER tree (Turakhia et al., 2021). However, here we have also observed that the inclusion of a large diversity in the reference can lead to distributed abundance assignments between closely related (sub)-lineages, reducing the true relative abundance of a lineage (***Figure S5*** and ***Figure S6***). Of course, the impact can be reduced by limiting lineage coverage to a specific time period, but this, in turn, can also affect frequency assignments.

In more detail, both approaches performed similarly in detecting and estimating spike-in lineage abundances for the *Standards* data set ***Figure 2***. The predictions are more similar on the parent-lineage level compared to the sub-lineage level. If their estimations differ, this can be mostly attributed to differences in the mutations/lineages included in the respective reference data: for both approaches, the final predictions heavily depend on the construction of the reference data set. In addition, both approaches had diZculties differentiating closely related sub-lineages correctly.

For the *Pan-EU-GER* data set, both approaches reflect well the pandemic background in Germany during the time of sampling, but we detected some limitations and potential sources for bias in their general behavior. Again, the choice of marker mutations and reference lineages impacts the level of detection: lineage vs. sub-lineage level estimations, but also the amount of low abundance detection. Potentially, everything that is defined in the reference data set can be also detected, which might lead to an increased number of false positive predictions. The whole-genome sequences or mutations used to create the reference index impact the degree of ambiguity and, thus, (low abundant) false positive detection. This may explain why here both approaches predicted distinctly different abundances on the parent-lineage level compared to the other two benchmark experiments. Therefore, we think that especially the *sequence-based* approach requires the definition of a false positive threshold to differentiate between low abundant false positive hits and low abundant true positives.

For the *FFM-Airport* data set, both approaches detect also low-frequency lineages. Again, the *sequence-based* approach detects a distinctly higher amount of low abundant lineages, also reflecting the higher diversity of the reference index.

We performed an additional parameter benchmark to identify important key parameters impacting the *sequence-based* pseudo-alignment approach using VLQ-nf. One parameter having a strong effect on the results, is the alternative allele frequency (AAF) cutoff. In connection with the reference size (the number of genomes), we observed different effects of changing the AAF. Our experiments also showed that the effect of the same parameter changes (increasing or decreasing AAF) does not yield consistent results among the different data sets. The degree of lineage ambiguity depends on the considered composition of lineages and sub-lineages. The effect of included/excluded mutations due to adjusted AAF parameter settings is variable, as different mutations have different effects in differentiating lineages. The effect of those parameter changes is most notable among lineages that are more similar. We also observed that with a larger reference size, the effect of the AAF parameter becomes smaller and overall abundance estimations improve. Increasing the reference size implicitly adds low-frequency mutations into a lineage reference set which in part reduces the effect of increasing/decreasing the AAF threshold when selecting sequences. Additionally, depending on their phylogenetic impact, low-frequency mutations might help better differentiate lineages.

## Potential implications

Most importantly, we only selected two exemplary implementations of the *mutation-* and *sequence-based* approaches MAMUSS and VLQ-nf, respectively, out of an increasing number of scripts, tools, and pipelines becoming available for computational SARS-CoV-2 lineage estimation from wastewater sequencing (***Table 1***) (Karthikeyan et al., 2022; Pechlivanis et al., 2022; Valieris et al., 2022; Ellmen et al., 2021; Amman et al., 2022; Barker et al., 2021; Schumann et al., 2022; Gregory et al., 2021; Jahn et al., 2022; Gafurov et al., 2022; Posada-Céspedes et al., 2021; Baaijens et al., 2022; Korobeynikov, 2022; Kayikcioglu et al., 2023). Thus, our benchmark results also reflect and are limited by the individual characteristics of these two implementations. However, we focused on these two approaches to investigate the impact of reference design using implementations where we could easily control parameters and input. Currently, a comprehensive benchmark comparison for the existing SARS-CoV-2 wastewater analysis tools is lacking. The developers of Freyja compared a selection of tools on a spike-in mixed sample (Karthikeyan et al., 2022) where they found that Freyja outperformed VLQ (Baaijens et al., 2022) in accuracy at higher expected proportions and observed noticeably longer computation times for both VLQ and LCS (Valieris et al., 2022). To counteract the effect on lineage abundance detection, some methods filter the mutations considered for lineage assignment based on sequencing depth (Amman et al., 2022) or adjust their mathematical model for differences in depth and coverage and expected error rates (Karthikeyan et al., 2022; Gafurov et al., 2022). In a similar context, the PiGx tool addresses the limitations of estimating lineages at low abundances by weighting specific signature mutations for lineages that are expected to occur at low frequencies (Schumann et al., 2022). As a next step, a broader evaluation of all available tools for the analysis of SARS-CoV-2 wastewater sequencing data is urgently needed to guide usage and further development.

## Conclusion

Academic researchers have pioneered wastewater monitoring of SARS-CoV-2 and overcome several technical and methodological challenges (Hoar et al., 2022). Thanks to these efforts, wastewater-based pathogen surveillance has rapidly become a valuable public health tool for detecting SARS-CoV-2 that can excellently complement syndromic surveillance or other monitoring tools. However, public health authorities are now faced with the task of integrating these achievements into robust and continuous public health surveillance systems that can be operated and expanded over the long term. For the inclusion of wastewater-based pathogen surveillance data, performance parameters must be defined and communicated to the public health authorities. In this context, continuous updating of reference data sets, in the context of retrospective analyses or time series, is essential to ensure comparability between time points. Especially for continuous sampling and analysis of wastewater-based SARS-CoV-2 sequencing data, the reference design must also be adjusted. Otherwise, a lineage defined with a delay might be present in older samples but was not detected only because it was not part of the reference at that time. However, harmonizing the reference used would require recalculating older abundance estimates, which may conflict with the standard reporting requirements of public health authorities. However, this problem is not specific to wastewater-based SARS-CoV-2 sequencing data, but also applies to genomics sequencing of patient samples. One solution might be to focus not only on lineages, but also to report mutations that are not affected by any nomenclature scheme and are not subject to delayed definitions. On the other hand, it is undeniable that lineages played a crucial role in communication during the COVID-19 pandemic.

The detection of *cryptic* (novel, undescribed) virus variants in wastewater samples is another powerful application for wastewater sequencing data, especially when clinical testing capabilities and monitoring systems are limited or reduced. In this context, approaches utilizing artificial intelligence might present a promising next step for the improved detection of cryptic SARS-CoV-2 lineages from wastewater sequencing data and potential outbreaks, although right now not much in use (Abdeldayem et al., 2022). Finally, the lessons learned from the sequencing efforts and implementations for SARS-CoV-2 detection from wastewater sequencing data can and should be adapted to other pathogens in the future.

## Methods

### Benchmark data set #1: Synthetic mixture ***Standards***

We procured synthetic SARS-CoV-2 RNA samples (Twist Biosciences), which were used to prepare 16 different mixtures (***Table 2***) containing different SARS-CoV-2 variants. From the pooled RNA, cDNA was synthesized using SuperScript™ VILO™ Master Mix (Thermofisher Scientific), followed by library preparation using the Ion AmpliSeq SARS-CoV-2 Research Panel (Thermofisher Scientific) according to the manufacturer’s instructions. This panel consists of 237 primer pairs, resulting in an amplicon length range of 125–275 bp, which cover the near-full genome of SARS-CoV-2. We performed multiple sequencing runs to achieve at least 1 million mapped reads per sample. For each sequencing run, eight libraries were multiplexed and sequenced using an Ion Torrent 530 chip on an Ion S5 sequencer (Thermofisher Scientific) according to the manufacturer’s instructions. The raw sequence data were uploaded to the National Center for Biotechnology Information (NCBI) Sequence Read Archive (SRA) under BioProject number PRJNA912560.

### Benchmark data set #2: ***Pan-EU-GER***

We obtained seven real samples from March 2021 from a large European study and collected in Germany (Agrawal et al., 2022b), mainly comprising the VOC Alpha (*Pan-EU-GER*, SRX11122519 and SRX11122521–SRX11122526).

### Benchmark data set #3: ***FFM-Airport***

We selected one sample from the end of 2021 including first signals of the VOC Omicron obtained from wastewater at the international airport in Frankfurt am Main, Germany (*FFM-Airport*, SRR17258654) (Agrawal et al., 2022a).

### Data processing: ***mutation-based*** reference design and lineage proportion estimation via MAMUSS

We used the SARS-CoV-2 Research Plug-in Package, which we installed in our Ion Torrent Suite software (v5.12.2) of Ion S5 sequence. We used the SARS_CoV_2_coverageAnalysis (v5.16) plugin, which maps the generated reads to a SARS-CoV-2 reference genome (Wuhan-Hu-1-NC_045512/MN908947.3), using TMAP software included in the Torrent Suite. The summary of mapping of each sample mentioned in ***Table 2*** is provided in ***Table S2***. For mutation calls, additional Ion Torrent plugins were used as described previously (Agrawal et al., 2023). First, all single nucleotide variants were called using Variant Caller (v5.12.0.4) with “Generic - S5/S5XL (510/520/530) - Somatic - Low Stringency” default parameters. Then, for annotation and determination of the base substitution effect, we used COVID19AnnotateSnpEff (v1.3.0.2), a plugin developed explicitly for SARS-CoV-2 and based on the original SnpEff (Cingolani et al., 2012). To construct reference marker mutation sets for MAMUSS, we used data from GISAID (https://www.gisaid.org) (Shu and McCauley, 2017) to reconstruct reference mutation profiles for each virus variant. The lineage abundance estimation is based on the read depth and allele frequency of each mutation detected in a wastewater sample followed by a two-indicator classification and comparison to the pre-selected marker mutations characteristic for certain lineages. For further details see github.com/lifehashopes/MAMUSS.

### Data processing: ***sequence-based*** reference design and lineage proportion estimation via VLQ-nf

Instead of only relying on manually or algorithmically selected marker mutations, another computational approach utilizes, in a first step, full genome information. For example, Baaijens, Zulli, and Ott *et al*. presented a method to estimate the abundance of variants in wastewater samples based on well-established computational techniques initially used for RNA-Seq quantification (Baaijens et al., 2022). Here, the main idea is that quantification of different transcripts derived from the same gene is computationally similar to the abundance estimation of different SARS-CoV-2 lineages derived from the same parental genome. Via Kallisto (Bray et al., 2016), they perform pseudo-alignments of the raw reads against an index of pre-selected and down-sampled full genome SARS-CoV-2 sequences with respective lineage information. Therefore, their approach may be less influenced by the pre-selection of mutations based on clinical relevance, frequency, or other parameters that mostly drive *mutation-based* tools, and thus may be better suited for sub-lineage discrimination. The approach comprises two steps: 1) selection of reference genome sequences for index construction and 2) pseudo-alignment of the reads and lineage abundance estimation. First, a reference data set of SARS-CoV-2 genome sequences must be selected. For that, we use data from GISAID (Shu and McCauley, 2017) and filter for human-host sequences, N-count information, pangolin annotation (Rambaut et al., 2020; O’Toole et al., 2021), origin (country, continent), and sampling date. This metadata is used to pre-select sequences based on geographic origin (continent, country), a sampling time frame, and the number of N bases. Next, the pipeline performs a variant calling against a reference sequence (per default index Wuhan-Hu-1, NC_045512.2) and subsequently samples sequences to select characteristic mutation profiles for each input lineage. Within a lineage, sequences are sampled based on an alternative allele frequency cutoff (e.g., AAF>0.5) so that each mutation is represented at least once until an upper limit of sequences per lineage is reached. From this downsampled and representative set of full genome sequences, a Kallisto index is constructed. Now, the raw reads from a FASTQ 1le are pseudo-aligned against this index and lineage abundances are estimated similarly to the estimation of transcript abundances in an RNA-Seq scenario.

For our comparative study, we used the initial idea and code base from Baaijens, Zulli, and Ott *et al*.(Baaijens et al., 2022) (https://github.com/baymlab/wastewater_analysis, version from September 16, 2021, with commit hash 61dd29df*) and implemented a Nextflow (Di Tommaso et al., 2017) pipeline (https://github.com/rki-mf1/VLQ-nf) with the purpose of automating the steps and making our analyses fully reproducible. In this context, we discovered some issues in the pipeline version 61dd29df* of Baaijens, Zulli, and Ott *et al*. and implemented minor adjustments. This includes updating data processing scripts according to the most recent GISAID data format and allowing the sequence selection based on alternate allele frequencies (AAF) to consider multi-allelic sites. Meanwhile, those issues have been addressed with similar code changes by the authors in their current pipeline version. Furthermore, we adjusted the AAF 1lter to 1rst sample sequences so that all passing mutations are included at least once before increasing the reference setup to the number of maximum sequences per lineage. In pipeline version 61dd29df*, sequences are selected for the reference index if they carry an AAF filter passing mutation that is not yet covered until the reference set for the respective lineage meets the maximum allowed number of sequences. We wondered if this sampling strategy might yield a reference index that does not suZciently represent the characteristic mutation profile of a lineage and could introduce potential sources of bias during reference reconstruction. We addressed this issue by implementing an AAF 1ltering that 1rst retrieves a minimal reference database for every lineage such that all AAF filter passing mutations are captured at least once by as few sequences as possible. In a second step, we incremented each lineage reference to match a maximum number of sequences using the remaining sequences passing the filter. We ran our pipeline version v1.0.0 for all analyses in this benchmark study.

### Reconstruction of indices for the ***sequence-based*** approach

The *sequence-based* (VLQ-nf) approach highly depends on the selection and reconstruction of the reference data set for the Kallisto index. Thus, we reconstructed different indices for our three benchmark data sets to mimic the pandemic situation during the time of sampling. For all indices, we used GISAID data and extracted subsets based on metadata filters.

For the benchmark of the 16 mixed *Standards*, we constructed a reference data set comprising the included SARS-CoV-2 lineages. We selected a time frame of two weeks around the peak of global incidences (based on outbreak.info (Gangavarapu et al., 2023), accessed April 1st, 2022) for each lineage included in the mix (***Table 3***). We only kept records with at least 29,500 non-ambiguous bases. Because we also included the original Wuhan-Hu-1 reference sequence in mixed samples Mix_01-Mix_05 and Mix_08, we 1rst excluded all A.1 sequences from the preselected set. Then, we selected reference sequences with characteristic mutation profiles for all lineages except A.1 as described before allowing a maximum number of 1ve sequences per lineage. Now, we added the sampled A.1 sequences again to the final reference set manually. Otherwise, the A.1 sequences would have been excluded by the pipeline because they don’t show any AAF in comparison to the Wuhan-Hu-1 reference. On average, we selected 1ve sequences for a lineage to capture every mutation against the wildtype with an AAF>0.25 (within-lineage variation) and a maximum of 1ve allowed sequences per lineage.

**Table 3.** For each lineage in the *Standards* data set, we selected the time frame where infection numbers peaked globally according to outbreak.info (Gangavarapu et al., 2023). Based on the listed time frames, we sampled genome sequences from GISAID for reference reconstruction. We downloaded the GISAID records on 02 March 2022.

| Lineage | Time frame |
| --- | --- |
| A.1 | 2020-03-01:2020-03-14 |
| B.1.1.7 | 2021-05-01:2021-05-14 |
| B.1.351 | 2021-01-20:2021-02-02 |
| P.1 | 2021-04-20:2021-05-03 |
| B.1.526 | 2021-03-20:2021-04-02 |
| BA.2 | 2022-02-01:2022-02-14 |
| BA.1 | 2021-12-01:2021-12-14 |
| B.1.617.2 | 2021-06-25:2021-07-08 |
| AY.1 | 2021-08-01:2021-08-14 |
| AY.2 | 2021-06-25:2021-07-08 |

For the *Pan-EU-GER* samples (collected between 10th and 30th March 2021), we reconstructed the reference from GISAID records we downloaded on 27 January 2022. We selected only European sequences sampled between February 1st, 2021, and April 30nd, 2021, with at least 29,500 non-ambiguous bases. We did not only select sequences from Germany to also mimic variant influx from other European countries. On average, we then selected three sequences per lineage to capture every mutation against the wildtype with an AAF>0.25 (within-lineage variation) and allowing at most 1ve reference sequences per lineage.

For the *FFM-Airport* data set, we reconstructed the reference from GISAID records we downloaded on 11 February 2022. We selected sequences from European and South African samples sampled between October 1st, 2021, and December 31st, 2021, again with at least 29,500 non-ambiguous bases. On average, four sequences were selected for a lineage to capture every mutation against the wildtype with an AAF>0.25 (within-lineage variation). Again, we allowed at most 1ve sequences to be included per lineage.

### Lineage-abundance estimation with the ***sequence-based*** approach

After reconstructing different reference indices for our benchmark data sets, we used specific Kallisto commands implemented in a Nextflow pipeline to prepare Kallisto mapping indices, compute pseudo-alignments of each benchmark data set against its reference index, and estimate lineage abundances following the original idea and code of Baaijens, Zulli, and Ott *et al*.(Baaijens et al., 2022).

First, we built a Kallisto index from the reference database (default k-mer=31). Next, for each sample in a benchmark data set, we pseudo-aligned all reads against the corresponding Kallisto index and estimated the abundance of each reference sequence in the sample. We quantified our benchmark data sets in single reads mode with an average fragment length of 200 nt with a standard deviation of 20 nt. Finally, a customized script groups the estimated abundances by the lineage annotation of the respective sequences and sums them up into a final lineage abundance estimation for the analyzed sample. For the *Pan-EU-GER* and *FFM-Airport* data sets, we further summarized the estimated abundances by the country information of the analyzed samples to compare the pseudo-alignment and *mutation-based* approach on the country level.

### Assessing parameter impact and potential bias with the pseudo-alignment approach

We performed parameter escalation experiments with our three benchmark data sets using the *sequence-based* method (VLQ-nf) to assess the impact of the AAF threshold and the cutoff for a maximum number of sequences per lineage on lineage abundance estimation. More importantly, we used the resulting observations to inform our choice of parameters used for the 1nal bench- marking against the *mutation-based* method (MAMUSS). In this context, we aimed at determining a setting with a good prediction performance and reasonable computational effort without manipulating the benchmark in favor of the *sequence-based* method. For every benchmark data set, we constructed reference indices over a range of 12 possible parameter combinations. For the AAF threshold, we iterated over [0.25, 0.5, 0.85] to cover lower, medium, and high threshold values to define the characteristic mutation profiles. For the maximum number of sequences per lineage, we built the reference index using the minimal sequence sets possible, 5, 10, and 20 sequences per lineage. After lineage abundance estimation with each reference index on the *Standards* data set, we evaluated prediction performance based on the ground truth lineage abundances. For the *FFM-Airport* and *Pan-EU-GER* data, we assessed prediction performance by comparing estimated lineage abundances with the pandemic background at the respective time and location.

### Reproducibility of the pseudo-alignment approach

Our Next2ow pipeline of the pseudo-alignment approach freely available at github.com/rki-mf1/VLQ- nf generates the reference database in the format of a CSV 1le containing the metadata information of the 1nal Kallisto index and a FASTA 1le containing the corresponding sequence data. In the current version v1.0.0, the reference CSV and FASTA can be exactly replicated using the same input data resource and index reconstruction parameters which leads to slightly different results at every analysis run. The reference CSV is not reproducible due to misplaced random sampling seeds and a missing record sorting strategy in the AAF-based sequence filtering step during reference reconstruction.

However, given already reconstructed reference indices (final CSV and FASTA reference) or an already built Kallisto index, lineage detection and quantification are deterministic. We deposit our used Kallisto indices at osf.io/upbqj.

### Availability of source code and requirements

Here, we provide the specifications of our Nextflow implementation (VLQ-nf) of the *sequence-based* approach originally presented by Baaijens, Zulli, and Ott *et al*.(Baaijens et al., 2022) and the code for the *mutation-based* approach, MAMUSS.

- Project name: VLQ-nf
- Project home page: https://github.com/rki-mf1/VLQ-nf
- Operating system(s): Linux, Mac, Windows via Linux subshell
- Programming language: Next2ow
- Other requirements: Conda
- License: GPL-3.0
- Project name: MAMUSS
- Project home page: https://github.com/lifehashopes/MAMUSS
- Operating system(s): Linux, Mac
- Programming language: R
- Other requirements: R packages are listed in the repository
- License: CC0 1.0 Universal

### Availability of supporting data and materials

The data sets supporting the results of this article are available in the Open Science Framework repository, osf.io/upbqj.

## Declarations

### List of abbreviations

- AAF - alternative allele frequency
- FFM-Airport - one sample from the end of 2021 including first signals of the VOC Omicron obtained from wastewater at the international airport in Frankfurt am Main, Germany (Agrawal et al., 2022a)
- MAMUSS - *mutation-based* approach for SARS-CoV-2 lineage abundance estimation
- Pan-EU-GER - seven samples from early 2021 from a large European study and collected in Germany, mainly comprising the VOC Alpha (Agrawal et al., 2022b)
- Standards - synthetic scenario of 16 “spike-in” mixture SARS-CoV-2 samples
- VLQ-nf - *sequence-based* approach for SARS-CoV-2 lineage abundance estimation, inspired by the original VLQ (Baaijens et al., 2022)
- WBE - wastewater-based epidemiology

### Ethical Approval (optional)

Not applicable.

### Consent for publication

Not applicable.

## Funding

### Author***’***s Contributions

SA, SL, and MH provided conceptualization and study design. SA implemented the MAMUSS approach and analyzed corresponding data. EA implemented the VLQ-nf approach and analyzed corresponding data. SA and LO conducted wet lab experiments to generate and sequence synthetic mixtures. EA, SA, and MH performed the computational comparisons and generated the figures. All authors actively participated in the writing and editing of the manuscript. All authors have read and agreed to the published version of the manuscript.

## Supplement

### Alternative allele frequency and size of reference database impact the ***sequence- based*** method but the effects are dependent on sample composition

#### Pan-EU-GER

Across all samples and experiments, we found the predictions of the *sequence-based* method to re2ect the pandemic background in Germany well. Alpha and its sub-lineages were among the most prominent predictions within the time frame of wastewater sampling (Supplementary ***Figure S3***). For most samples, we found Alpha and Q.1 to be the most abundant (sub-)lineages. The *sequence-based* method predicted distinctly varying abundances for sub-lineages other than Alpha, Beta, Gamma, or B.1.617 (summarized as “Other”) across the *Pan-EU-GER* samples. We chose a cutoff of 1 % abundance to differentiate true positive predicted lineages from false positive noise. On average, we found the pseudo-alignment-based approach to detect around 20–30 % abundance of noise across all samples and parameter settings. At the minimum reference size (Supplementary ***Table S1***), we observed for some samples a slightly decreasing amount of noise and a slightly increasing abundance for Alpha sub-lineages and “Others” when increasing the AAF threshold (e.g., sample INF_21051_D). We found, that the number of “Others” sub-lineages above 3 % abundance decreased with increasing reference size. Across all experiments, we found the sample INF_21011_D to be the only one to be predicted with one or two “Others” sub-lineages of at least 3 % abundance.

With increasing AAF threshold, we found distinct shifts in the estimated abundances for B.1.1.7 and Q.1. We observed those shifts to behave complementary but not consistently across all reference sizes: At the minimum reference size, we observed Alpha abundance predictions to distinctly increase and Q.1 abundances to decrease across all samples with increasing AAF threshold. Conversely, for reference size 5, we found Alpha abundance predictions to first increase and then decrease again with increasing AAF threshold. Vice versa, we observed Q.1 abundances to decrease and then increase again. At the largest reference size of 20 sequences per lineage, we observed a consistent decrease in Alpha abundance estimates and a consistent increase in Q.1 abundance estimates with increasing AAF threshold. Furthermore, we found abundances of other Alpha sub- lineages like Q.4 and Q.6 to also increase and decrease across varying parameter settings without following a clear pattern, but found the predicted abundances to not change as distinctly.

Overall, we found the performance of the *sequence-based* method to be mostly robust with varying settings for the AAF threshold and reference size. We observed the impact of those parameter changes to be stronger for more closely related lineages in a sample and in some cases to become weaker at larger reference sizes.

#### FFM-Airport

Across all parameter settings, the resulting abundance profiles for the *FFM-Airport* data set re2ected the pandemic background in Europe and South Africa well around the time frame of wastewater sampling: the *sequence-based* method estimated Delta and its sub-lineages to represent the most abundant lineages and detected small proportions of Omicron (Supplementary ***Figure S4***). We chose a cutoff of 1 % abundance to differentiate true positive lineages from false positive noise and labelled sub-lineages with a minimum abundance of 3 %. Because the *sequence-based* method detected Omicron sub-lineages at abundances below 3 %, the quantified levels are not labelled and due to the scale of Supplementary ***Figure S4*** not visible. However, when grouped by parent lineage, the predicted Omicron proportions become obvious. On average, we found the *sequence-based* method to detect around 50 % abundance of noise across all parameter settings.

At the minimum reference size (Supplementary***Table S1***), we observed a decreasing amount of low abundant noise with increasing AAF threshold. In contrast, with larger reference sizes, we found the amount of low abundant noise to change slightly and not follow a consistent pattern. Overall, we found the amount of noise to increase with increasing reference size. We observed the abundance estimates to increase for individual Delta sub-lineages with increasing AAF threshold. Specifically, we found the set of the most abundant Delta sub-lineages to change at every increase. Some examples for Delta sub-lineages that alternately were estimated among the most abundant lineages within a sample are AY.43.1 and AY.43.2, AY.43.3 and AY.42, and AY.121 and AY.122. When considering the lineage abundance profiles grouped by parent lineages, we found the predicted abundance profiles to not change distinctly across different parameter settings(Supplementary ***Figure S4***.

Finally, we found different settings for the AAF threshold and reference size to not distinctly affect the performance of the *sequence-based* method. We observed variations in the abundance estimates among multiple Delta sub-lineages.

**Supplementary Figure S1.**
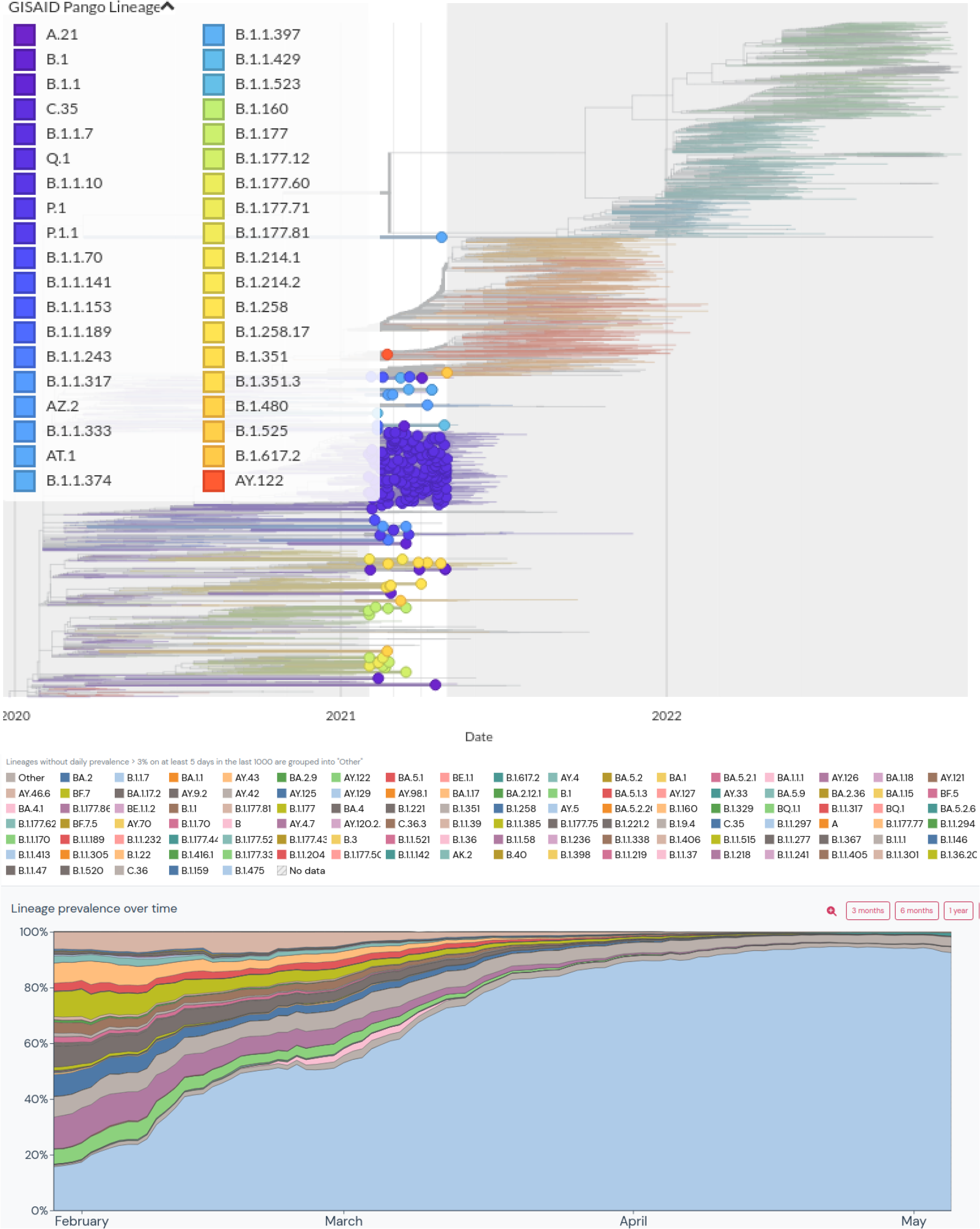
Top: The pandemic background across Europe between 01 February and 30 April 2021 was built with Nextstrain.org. Bottom: The outbreak.org variant report for Germany displaying the SARS-CoV-2 lineage prevalence from February to March 2021 based on GISAID sequence data. The most dominant lineages in the plot from bottom to top: light blue = B.1.1.7, light green = B.1, purple = B.1.177.86, light grey = other, blue = B.1.258, dark grey = B.1.221, yellow = B.1.177, orange = B.1.160, light brown = B.1.177.81

**Supplementary Figure S2.**
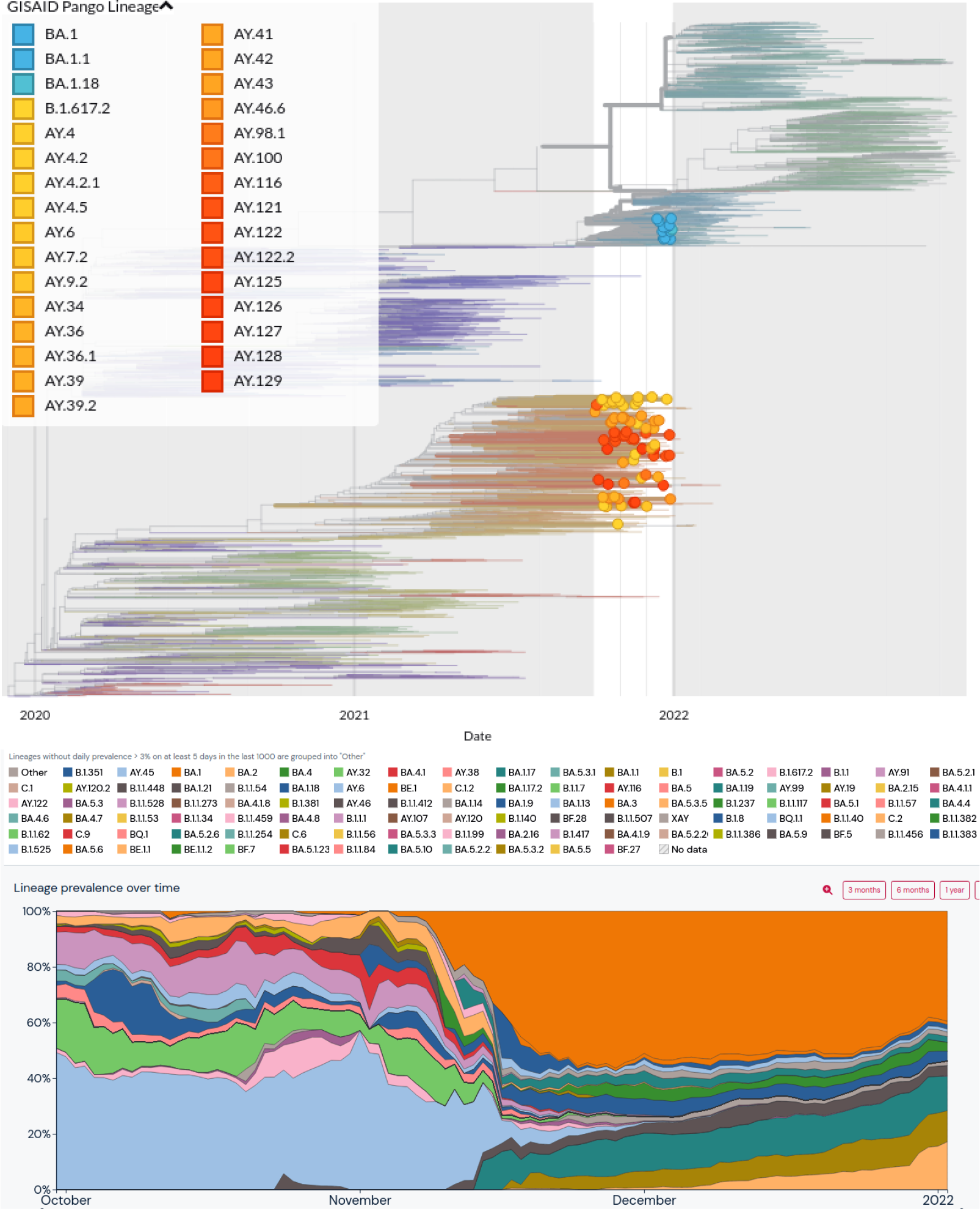
Top: The pandemic background across Europe between 01 October and 31 December 2021 was built with Nextstrain.org. Bottom: The outbreak.org variant report for South Africa displaying the SARS-CoV-2 lineage prevalence from October to December 2021 based on GISAID sequence data. The most dominant lineages comprise sub-lineages of Delta and Omicron, but also B.1.351 (light green) and C.2 (light orange).

**Supplementary Figure S3.**
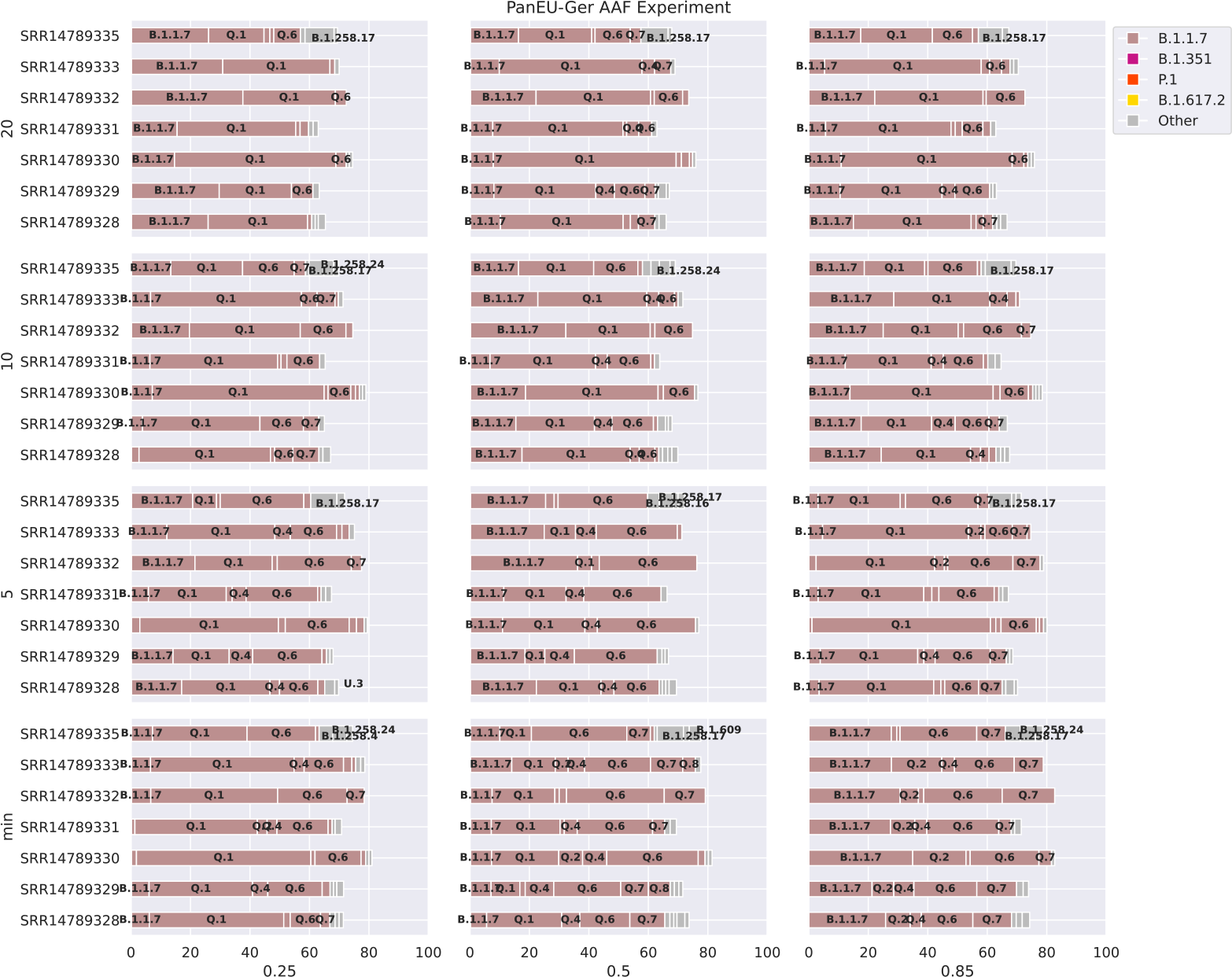
Results for the parameter escalation experiments on the *Pan-EU-GER* samples using the *sequence-based* method using pseudo-alignment implementation. We analyzed the data set with different parameterization for reference construction (x-axis: increasing AAF threshold, y-axis: increasing maximum number of sequences per lineage). Abundance predictions are displayed at a minimum threshold of 1 % and labelled at a threshold of 3 %. When comparing with the pandemic background at the time of wastewater sampling, we observed the AAF threshold and the maximum number of sequences per lineage to impact the abundance proportions among Alpha and Q.1 the most. With more sequences per lineage in the reference, we found the impact of the AAF filter on the observed ambiguities to decrease. We found significantly more low abundant sub-lineages predicted in the real wastewater data compared to the *Standards* data set and found those low abundant predictions to mostly not change distinctly across varying parameterization.

**Supplementary Figure S4.**
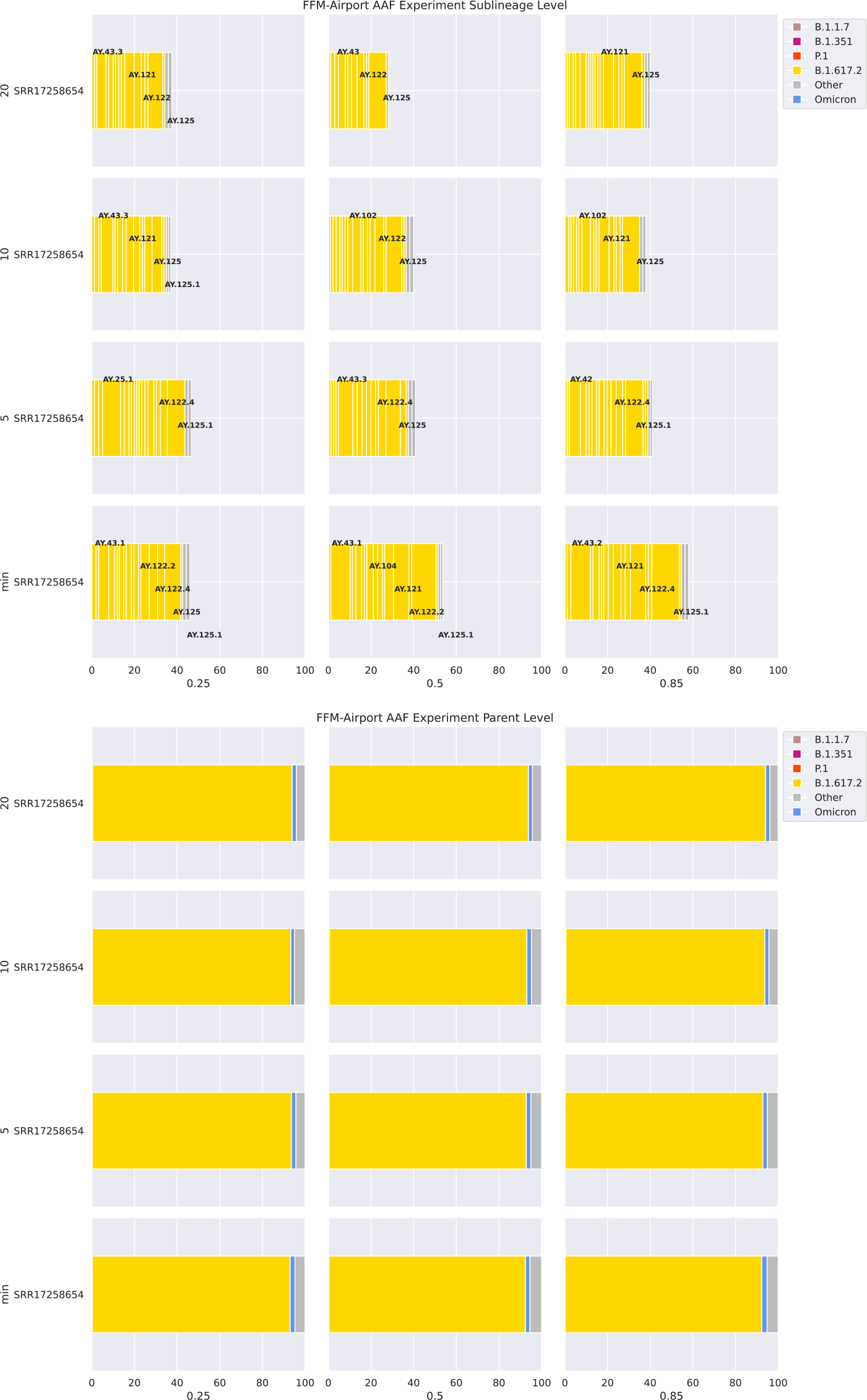
Results for the parameter escalation experiments on the *FFM-Airport* data set using the *sequence-based* method. We analyzed the data set with different parameterization for reference construction (x-axis: increasing AAF threshold, y-axis: increasing maximum number of sequences per lineage). Top: Abundance predictions are displayed at a minimum threshold of 1 % abundance and labelled at a threshold of 3 % abundance. When comparing with the pandemic background at the time of wastewater sampling, we observed the following: Overall, we found more sub-lineages predicted with abundance below 1 % compared with the *Standards* data set and the *Pan-EU-GER* set. The *sequence-based* method detected more low abundant sub-lineages with increasing reference size and slightly less low abundant sub-lineages with increasing AAF threshold. Both the AAF threshold and the reference size showed to impact lineage ambiguities among Delta sub-lineages. Bottom: All abundance predictions are displayed as grouped by their parent lineage. We did not 1nd the abundance predictions for parent lineages to change distinctly across experiments.

**Supplementary Figure S5.**
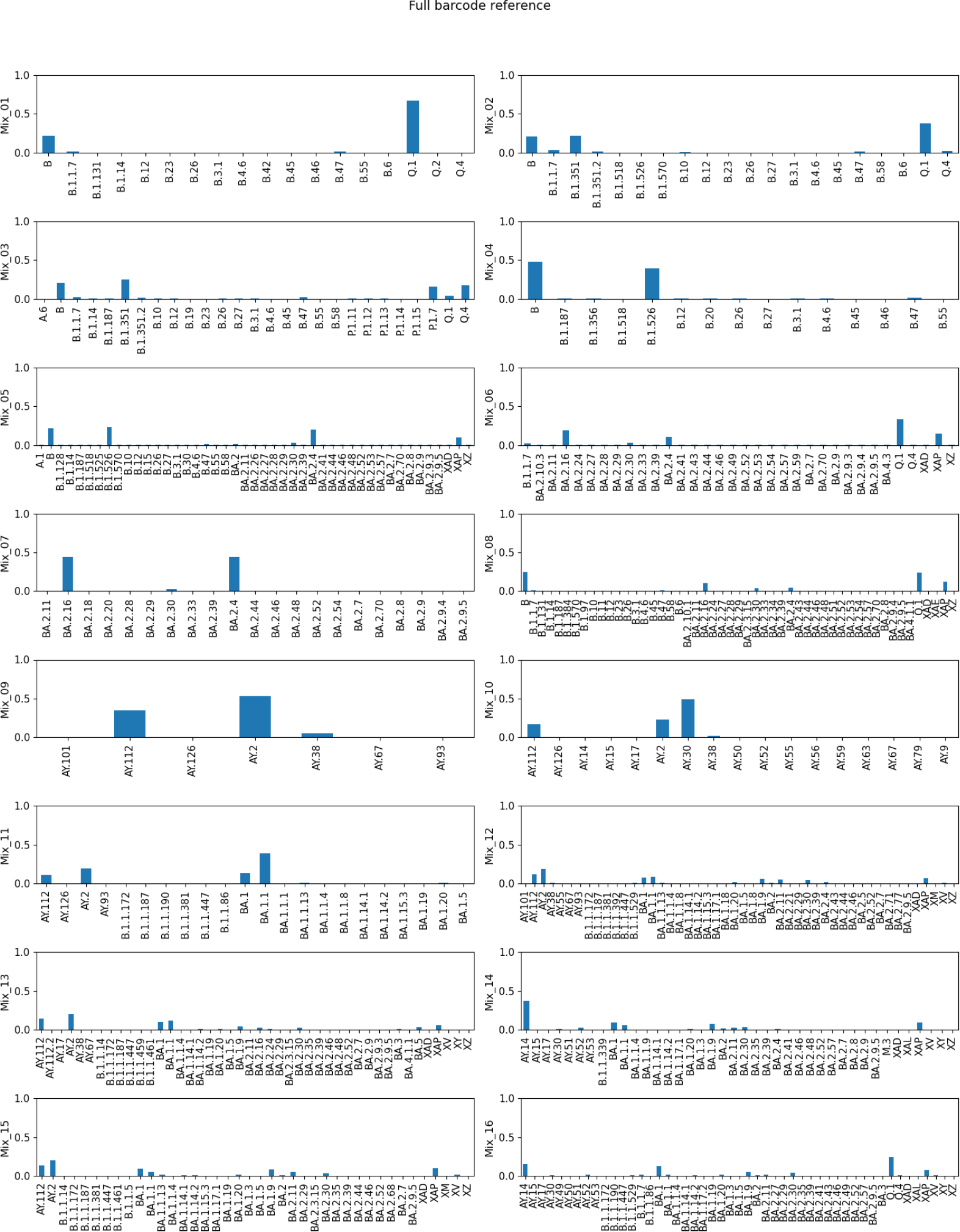
SARS-CoV-2 lineage abundance assignments via Freyja Karthikeyan et al., 2022 (v1.3.12) for the *Standards*. We used the full reference UShER set as provided as a default by the tool. In this case, multiple sub-lineages were predicted and frequencies were distributed among them, resulting in a reduced frequency estimate for the true (parental) lineage and an increase in low-frequency detections. For example, in Mix_07 the sub-lineages BA.2.16 and BA.2.4 were predicted with almost 50 %, respectively, while the included lineage BA.2 was not assigned (compare ***Figure S6***).

**Supplementary Figure S6.**
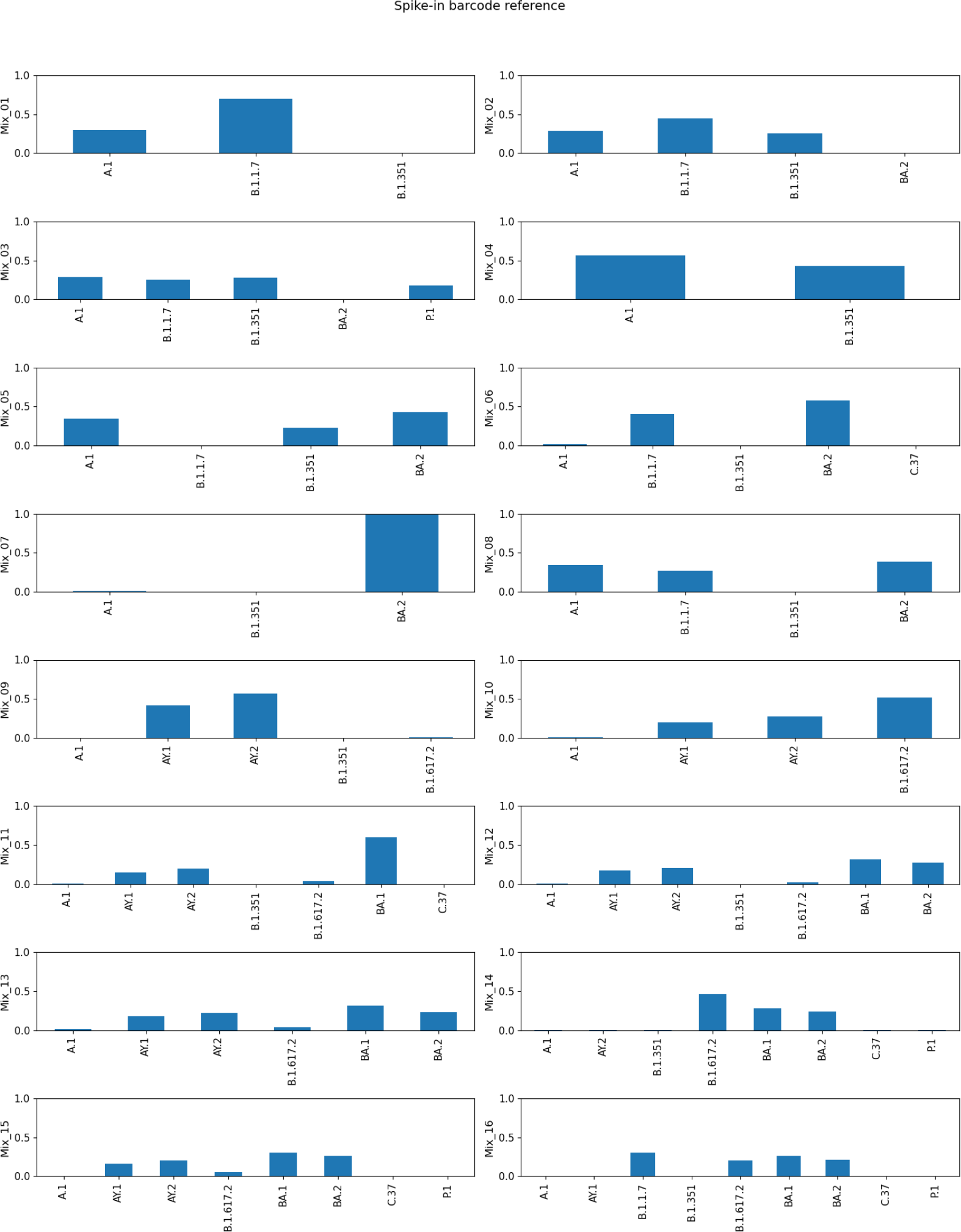
SARS-CoV-2 lineage abundance assignments via Freyja Karthikeyan et al., 2022 (v1.3.12) for the *Standards*. We reduced the reference UShER set to the lineages part of our artificial mixtures, instead of using the full UShER barcode data set as shown in ***Figure S5***.

**Supplementary Table S1.** Table showing the minimum reference sizes across the different alternative allele frequency (AAF) thresholds considered in the parameter escalation experiments across our three benchmark data sets. Here, we list the minimum number of genome sequences required per lineage to capture every mutation with an AAF above the considered AAF threshold at least once based on the implemented sampling strategy during reference construction. The *Standards* reference database required the largest number of sequences to capture the predefined genomic variation. Overall, we observed that with an increasing AAF threshold, the minimum reference sizes per lineage decreased across all three benchmark data sets.

| AAF threshold | Minimum number of sequences per lineage |  |  |
| --- | --- | --- | --- |
|  | Standards | Pan-EU-Ger | FFM-Airport |
| 0.25 | 20 | 3 | 3 |
| 0.5 | 10 | 2 | 2 |
| 0.85 | 10 | 1 | 1 |

**Supplementary Table S2.** Table summarizing the mapping of each *Standards* sample.

| **SampleID** | **Total number of reads** | **Number of mapped reads** | **Average target base coverage depth\*** |
| --- | --- | --- | --- |
| Mix_01 | 11585401 | 11409164 | 67129 |
| Mix_02 | 8000781 | 7872877 | 45648 |
| Mix_03 | 7182909 | 7082168 | 41674 |
| Mix_04 | 11156327 | 11033124 | 65477 |
| Mix_05 | 9509819 | 9358747 | 54475 |
| Mix_06 | 12228003 | 12005442 | 69829 |
| Mix_07 | 11227258 | 10999971 | 63366 |
| Mix_08 | 7991289 | 7886999 | 45904 |
| Mix_09 | 22189148 | 21855903 | 21855903 |
| Mix_10 | 8685555 | 8614170 | 8614170 |
| Mix_11 | 2564976 | 2535182 | 2535182 |
| Mix_12 | 2581594 | 2557744 | 2557744 |
| Mix_13 | 8429568 | 8295217 | 8295217 |
| Mix_14 | 6246713 | 6173867 | 6173867 |
| Mix_15 | 11888445 | 11752739 | 11752739 |
| Mix_16 | 6139010 | 6092648 | 6092648 |

\*Target sequence was the SARS-CoV-2 reference genome (Wuhan-Hu-1)

